# The fungal transcription factor SmpR coordinates secondary metabolism and antibacterial defence in *Aspergillus fumigatus* during interspecies interaction

**DOI:** 10.64898/2026.04.05.716574

**Authors:** Sophie Tröger-Görler, Raghav Vij, Sascha Schäuble, Maira Rosin, Henrik Schweder, Peter Hortschansky, Volker Schroeckh, Amelia E. Barber, Olaf Kniemeyer, Sascha Brunke, Gianni Panagiotou, Bernhard Hube, Axel A. Brakhage

## Abstract

*Aspergillus fumigatus*, an opportunistic human fungal pathogen, encodes numerous secondary metabolite biosynthetic gene clusters (BGCs) that are tightly regulated and often remain silent under standard conditions. Co-cultivation with *Streptomyces rapamycinicus* or treatment with the secondary metabolite from this species, the arginoketide azalomycin F, induce the otherwise silent fumicycline (*fcc*) BGC of *A. fumigatus*. To elucidate the underlying regulatory circuitry, we performed transcriptome analyses of *A. fumigatus* exposed to azalomycin F or co-cultured with *S. rapamycinicus*. Both conditions triggered a coordinated antibacterial response, characterized by induction of specific secondary metabolites and antibacterial effectors, alongside repression of other BGCs, including those for fusarinine C, pyripyropene A, and fumagillin. Among the most strongly induced genes was a zinc cluster transcription factor, designated SmpR for <u>s</u>econdary metabolite <u>m</u>ultiple <u>p</u>athway regulator, which is conserved within Ascomycota. SmpR expression was selectively induced by azalomycin F, specific *Streptomyces* species and other bacteria isolated from soil such as *Kribbella* spp. and *Arthrobacter* spp.. Functional analyses revealed that SmpR is required for activation of the fumicycline BGC: its deletion reduced, whereas its overexpression enhanced fumicycline production independently of external stimuli. We further demonstrate that SmpR acts upstream of the pathway-specific regulator FccR and additionally controls multiple antibacterial BGCs, including those for hexadehydroastechrome, helvolic acid and xanthocillin. Together, our data identify SmpR as a key regulator coordinating antibacterial secondary metabolism in response to bacterial signals in *A. fumigatus*.

## INTRODUCTION

*Aspergillus fumigatus* is an opportunistic human pathogenic fungus known to cause severe infections like invasive aspergillosis particularly in immunocompromised patients (Morrissey *et al*., 2024). In 2022 it was added to the critical priority list of fungal pathogens by the world health organization (WHO, 2022). The fungus is known for its ability to produce a wide array of secondary metabolites (SMs) in axenic- or co-culture with other organisms (Yurchenko *et al*., 2023). Although not essential for *A. fumigatus* growth, these compounds play an important role in pathogenicity, defence and shaping it’s environment, and can be of applied interest (Boysen *et al*., 2021; Zhang *et al*., 2022; Brakhage, 2024). Genes involved in SM production are arranged in biosynthetic gene clusters (BGCs), which consist of several genes encoding proteins of different function in close proximity to each other (Keller, 2015). In *A. fumigatus,* more than 30 BGCs are known to produce SMs (Steenwyk *et al*., 2020). Most SMs can be assigned to the group of polyketides, terpenes, small peptides or hybrid molecules thereof (Keller, 2019).

In addition to genes encoding a synthase/synthetase and tailoring enzymes (for examples epimerases, p450 monooxygenases or hydroxylases), BGCs can encode pathway-specific transcription factors, however, this is not the case for all BGCs (Brakhage, 2013; Keller, 2019; Wang *et al*., 2021). Transcription factor genes not being part of BGCs are considered as global regulators that can coordinate the activation of BGCs with cellular responses (Macheleidt *et al*., 2016). In some cases, the cooperation of multiple transcription factors, encoded within or outside of BGCs, is required to ensure an adequate production of SMs (Ries *et al*., 2020; Paul *et al*., 2023).

BGCs are often silent under standard laboratory conditions, and only certain stimuli can activate their expression and consequently, the SM production. A stimulus activating at least some silent gene clusters relies on the co-incubation of a fungus, in this case different *Aspergillus* species, with bacteria such as soil-dwelling *Streptomyces* species. This results in the production of various SMs (Schroeckh *et al*., 2009; Rateb *et al*., 2013). Especially *S. rapamycinicus* is known for its capability to induce the formation of several SMs by *A. fumigatus* (König *et al*., 2013; Stroe *et al*., 2020). This co-cultivation strategy led to the identification of the *fcc* BGC (also called *nsc* BGC) in *A. fumigatus*, whose activation leads to the biosynthesis of the meroterpenoid fumicycline (also called neosartoricin) (König *et al*., 2013). The *fcc* cluster encodes a β-lactamase like thioesterase (*fccB*), a monooxygenase (*fccC*), a polyketide synthase (*fccA*), a tryptophan synthase (*fccD*) and an epimerase (*fccE*) gene and is regulated by the cluster-encoded transcription factor FccR (König *et al*., 2013; Chooi *et al*., 2013). Until now, there is little information about contributing factors enabling the activation of the *fcc* cluster gene (König *et al*., 2013). More insights into the regulatory and signaling circuits have been gained for the interaction of the filamentous fungus *Aspergillus nidulans* with *S. rapamycinicus* (Schroeckh *et al*., 2009). Here, the interaction results in the production of the fungal polyketide orsellinic acid (Nützmann *et al*., 2011). The inducing compound produced by the bacterium was found to be the polyketide azalomycin F which induces the otherwise silent fungal *ors* BGC (Krespach *et al*., 2023).

To shed light on the transcriptomic response of *A. fumigatus* to *S. rapamycinicus*, and its metabolite azalomycin F (AzF), we applied a RNA-seq analysis approach. As a result, we discovered the involvement of the Zn(II)_2_Cys_6_ transcription factor SmpR (secondary metabolite multiple pathway regulator) in fumicycline production. Most of the genes whose transcription was negatively affected by the deletion of *smpR* were associated with secondary metabolism. Notably all of the genes arranged in the *fcc* BGC (fumicycline production), parts of the *has* BGC encoding genes for production of hexadehydroastechrome (HAS), as well as the *hel* BGC and the *xan* BGC responsible for helvolic acid and xanthocillin production, respectively (Yin *et al*., 2013; Lv *et al*., 2017; Lim *et al*., 2018). We also showed a hierarchical relationship of SmpR with FccR. This finding suggests that SmpR acts as a more global regulator of the *fcc* BGC. Furthermore, SmpR may also play a role as a global regulator of the antibacterial response by activating antifungal proteins and peptides besides SMs in *A. fumigatus*.

## MATERIALS AND METHODS

### Strains and culture conditions

*A.fumigatus* strains were grown on *Aspergillus* minimal medium (AMM) agar plates (Brakhage and Van den Brulle, 1995) for 72 h at 37°C. Conidia were harvested in ultra-pure water and counted with a hemocytometer. Transformed *A. fumigatus* protoplasts were plated on AMM agar plates containing 100 µg/ml pyrithiamine, 150 µg/ml hygromycin or a combination of both. The *A. fumigatus* strains used in this study were either based on the *A. fumigatus* strain CEA17 11*KU80* (da Silva Ferreira *et al*., 2006) or the mutant strains Δ*smpR* (this study) and Δ*fccR* (König, 2015). All strains are listed in supplementary Table S1. Their construction and verification are reported in the supplementary material (Figure S1, Figure S2).

*Streptomyces* species used in this study are listed in Table S2. All used species, except *S. iranensis,* were cultured as described in Schroeckh *et al*., 2009. *S. iranensis* was grown for 96 h in tryptone soy broth (TSB) with 10 % (w/v) sucrose (Kieser *et al*., 2000) at 28°C, 180 rpm. For growth assessment of *Streptomyces* species with fumicyclines B and C (also known as neosartoricin B and neosartoricin C), species were grown in 48-well plates (Greiner bio-one, Frickenhausen, Germany) in the presence of 64 µg/ml fumicycline B or fumicycline C (Biomol, Hamburg, Germany), which were dissolved in methanol (MeOH). Growth was followed microscopically by a Keyence BX800 microscope (Keyence, Neu-Isenburg, Germany) after 24 h of incubation for a total of 96 h. Growth in wells with fumicyclines was compared to growth in wells without fumicyclines and wells containing MeOH.

Bacteria that were isolated from soil and used in this study are listed in Table S3. All soil isolates were grown as pre-cultures in TSB over night at 28°C, 180 rpm. For the inoculation of main cultures which were used for experiments, 50 µl of the pre-cultures were inoculated into fresh TSB and grown over night at 28°C, 180 rpm. *Bacillus subtilis* 168 and *Escherichia coli* DH5α were propagated in LB medium (Carl Roth, Karlsruhe, Germany) with shaking at 180 rpm at 28°C and 37°C, respectively.

### Isolation of soil bacteria

Soil samples were collected in Jena (Fürstenbrunnen, Jena, Germany) and given to 50 ml plastic tubes. The soil samples were then stored at −20°C until further use. After grinding the samples with a mortar and pestle, the soil samples were suspended in 9 ml of a Na_4_P_2_O_7_ solution with pH of 8. The suspension was shaken at room temperature for 30 minutes. After shaking, the soil was sedimented for 5 minutes. The supernatant was harvested and stored at 4°C. 9 ml of 0.5M NaCl were added to the sediment and ultrasonicated for 5 min at 50 W to separate remaining spores and organisms form the soil particles. The supernatant was again harvested. Serial dilutions (1:10, 1:100, 1:1.000, 1:10.000) of the supernatant were plated on AMM agar plates that were incubated at 28°C until colonies became visible. Identification of the isolated bacteria was achieved *via* 16S DNA amplification using PCR. For this reason, cultures of the bacteria were grown over night in at 28°C and shaken with 180 rpm in liquid TSB. Bacterial cells were pelleted via centrifugation with 14,000 rpm. The extraction of genomic DNA (gDNA) was carried out with the NucleoSpin® Microbial DNA Kit (Macherey Nagel, Düren, Germany). The PCR was conducted with primer pair OMK_505/OMK_506 (Table S4). PCR Products were loaded on a 1% agarose gel and run at 130 V for 25 minutes. The resulting bands were excised and purified via the Zymoclean™ Gel DNA Recovery Kit (Zymo Research, Freiburg, Germany) and sent for sequencing (Eurofins Genomics, Ebersberg, Germany). The species *Arthrobacter* spp. and *Kribbella* spp. were further analysed.

### Co-culture of bacterial strains with *A. fumigatus*

Co-cultivation of *Streptomyces* spp. with *A. fumigatus* strains was carried out as described before (Schroeckh *et al*., 2009). Briefly, fungal pre-cultures were grown in AMM for ∼16 h at 37°C, 200 rpm and filtered through 40 µm EASYstrainer™ (Greiner bio-one, Frickenhausen, Germany) and mycelium was transferred to fresh AMM. For strains with an inducible xylose-promoter, AMM with 1 % (w/v) glucose was substituted by AMM with 1 % (w/v) xylose. Fungal cultures were inoculated with 1/20 of the *Streptomyces* spp. cultures and co-cultured for 3 h for RNA extraction or 8 h for extraction of SMs. Cultures of the soil bacteria *Arthrobacter* spp., *Kribbella* spp. and the laboratory strains *B. subtilis* 168 and *E. coli* DH5α were conducted in AMM medium which was supplemented with 20 % (v/v) of TSB (for the soil isolates) or LB (for the laboratory strains) to promote bacterial growth. Cultures of *A. fumigatus* Δ*KU80* strain were prepared as described above. Bacteria were counted with a CASY^VIVO^ cell counter (OMNI life science, Bremen, Germany) and a defined inoculum of 1 x 10^6^/ml bacteria was added to the fungal culture. Samples for qRT-PCR were harvested after 3h of co-culture while cultures for SM extraction were harvested after 8h of co-culture. SMs were isolated as described below.

### Determination of growth curves

*Arthrobacter* spp. and *Kribbella* spp. were cultivated as described above. Experimental cultures were counted with a CASY^VIVO^ cell counter (OMNI life science, Bremen, Germany) and a defined inoculum of 5 x 10^5^ /ml bacteria in TSB was then added to the wells of a 96-well plate (Greiner bio-one, Frickenhausen, Germany) in the presence of different concentrations of pure fumicycline B or fumicycline C (Biomol, Hamburg, Germany). MeOH in different concentrations served as a growth control, the end concentration of MeOH did not exceed 1.3 % (v/v). The plates were incubated in a BioTek LogPhase 600 microbiology reader (Agilent, Santa Clara, United States) at 30°C, 500 rpm for 24 h. Measurement of optical density was carried out every 20 mins at 600 nm. All growth curves were determined in triplicates.

### RNA isolation and qRT-PCR

For transcriptomic analysis, three biological replicates of *A. fumigatus* Δ*smpR* and *A. fumigatus* CEA17 Δ*KU80* were grown in liquid culture as described above in the presence or absence of 10 µg/ml AzF which had been extracted from *S. iranensis* cultures (as described in Krespach *et al*., 2023) or co-cultured with *S. rapamycinicus* and harvested using Miracloth (Merck Millipore, Darmstadt, Germany). RNA isolation and DNase digestion were carried out as described in Abou-Kandil *et al*., 2024. RNA quality was analysed with the Qubit RNA Integrity and Quality (IQ) Assay kit (Thermo Fisher Scientific, Dreieich, Germany). The 260 nm/230 nm ratio was determined using a Nanodrop ND-1000 (Thermo Fisher Scientific, Dreieich, Germany) and total RNA was subjected to Illumina NovaSeq 6000 (Novogene, Cambridge, UK) sequencing yielding 20 million reads per sample.

For qRT-PCR analysis of target gene expression in *A. fumigatus*, strains were grown in the presence or absence of different bacteria. cDNA synthesis and qRT-PCR were carried out as described in Abou-Kandil *et al*., 2024. Target genes were normalized to the expression of the reference gene *act1* using the 2^-ΔΔCt^ method (Livak and Schmittgen, 2001). For qRT-PCR experiments involving soil bacteria, absolute quantification of mRNA concentrations was used, since the expression of target genes was below the detection threshold in the axenic *A. fumigatus* Δ*KU80* strain.

All used qRT-PCR primers had an efficiency of 90-110 %. Primers for target amplification, as well as their efficiency, are listed in Table S5 in the supplementary material.

### RNA-seq data processing

Preprocessing of raw reads including quality control and gene abundance estimation was achieved with the GEO2RNaseq pipeline (v0.100.3, Seelbinder *et al*., 2019) in R (version 3.6.3). For quality analysis FastQC (v0.11.5) was employed before and after trimming. Read-quality trimming was done with Trimmomatic (v0.36). Reads were rRNA-filtered using SortMeRNA (v2.1) with a single rRNA database combining all rRNA databases shipped with SortMeRNA. Reference annotation was created by extracting and combining exon features from corresponding annotation files. Reads were mapped against the reference genome of *A. fumigatus* Af293 (refSeq assembly ASM265v1) using HiSat2 (v2.1.0, paired-end mode). Gene abundance estimation was done with featureCounts (v2.0.1) in paired-end mode with default parameters. MultiQC version 1.7 was finally used to summarize and assess the quality of the output of FastQC, Trimmomatic, HiSat, featureCounts and SAMtools. The count matrix with gene abundance data without and with median-of-ratios normalization (MRN, Anders and Huber, 2010) were extracted. Raw files are accessible under the Gene Expression Omnibus accession number GSE288601. Differential gene expression was analysed using GEO2RNaseq. Pairwise tests were performed using four statistical tools (DESeq v1.38.0, DESeq2 v1.26.0, limma voom v3.42.2 and edgeR v3.28.1) to calculate *p* values and multiple testing corrected *p*-values using the false-discovery rate method q = FDR(p) for each tool. In addition, mean MRN, transcripts per kilobase million (TPKM) and reads per kilobase million (RPKM) values were computed per test per group including corresponding log_2_ of fold-changes. Gene expression differences were considered significant if they were reported to be significant by all four tools (q ≤ 0.05) and |log_2_(fold-change [MRN based]) | ≥ 1.

### Transcriptomic data visualisation and AlphaFold 3-based modelling

Kyoto encyclopedia of genes and genomes (KEGG) pathway and gene onthology (GO) term enrichment analyses were performed to identify significantly regulated pathways and genes. All analyses were conducted using R version 4.4.1. Genes that were significantly regulated by all four aforementioned statistical tools and exhibited a log_2_ fold change greater than 0.5 or less than −0.5 compared to the respective control were selected. Pathway enrichment analysis was performed using the clusterProfiler package (v4.4.2) in R (Wu *et al*., 2021). For GO term enrichment analysis, the enricher function was used with the following parameters: a minimum gene set size of 10, a maximum gene set size of 500, and a Benjamini-Hochberg adjusted *p*-value threshold of 0.05. Curated GO terms were downloaded from FungiDB (Alvarez-Jarreta *et al*., 2023). KEGG pathway enrichment analysis was conducted using the enrichKEGGfunction with the same parameters. Heatmaps for BGCs that were not annotated in GO or KEGG (fusarinine C, pyripyropene A, xanthocillin, helvolic acid, fumicyclines and hexadehydroastechrome) as well as genes encoding transcription factors, were created in R version 4.4.1 with the thresholds indicated in the figure legends. Only genes identified as significantly differently transcribed by DESeq, DESeq2, limma voom and edgeR were used.

For structure prediction of SmpR, the AlphaFold3 software was applied (Abramson *et al*., 2024). All gene annotations and protein sequences were retrieved from FungiDB (Alvarez-Jarreta *et al*., 2023).

### Isolation and LC-MS analysis of SMs

Extraction and detection of SMs were carried out as described previously (Stroe *et al*., 2020). For detection and analysis of natural products, 1 µL of the extracts was loaded onto a LUNAR Omega PS C18 LC-column with a particle size of 1.6 µm (Phenomenex, Aschaffenburg, Germany) coupled to a Vanquish Flex UHPLC system equipped with a PDA and an OrbitrapTM Exploris 120 mass spectrometer (Thermo Fisher Scientific, Dreieich, Germany). As a solvent system water/acetonitrile was used with 0.1 % (v/v) formic acid and a flow rate of 0.5 mL min^-1^ (program over 13 min: initial 5 % (v/v) acetonitrile hold for 0.5 min and increased to 100 % (v/v) over 8 min held at 100 % (v/v) for another 2 minutes and reversed to 5 % (v/v) over 2 min). Compounds were identified by comparison with authentic references (retention time, ultraviolet light spectrum, and HR-MS and MS/MS spectra). Identification of fumicyclines was achieved by comparison with the authentic reference neosartoricin B and neosartoricin C (Biomol, Hamburg, Germany).

The mass-to-charge ratios in this study are as follows: fumicycline A (*m/z* 423.5 [M-H]^-^), fumicycline B (*m/z* 441.5 [M-H]^-^), fumicycline C (*m/z* 483.5 [M-H]^-^) and helvolic acid (*m/z* 585.32 [M-H]^-^). Fumicycline B, fumicycline C and helvolic acid extracted ion chromatograms (EIC) were compared to EICs of the authentic standards (Figure S3).

### Luminescence measurement

The *A. fumigatus* strain *smpR-nluc* was cultured for 16 h in AMM at 37°C, 200 rpm, washed and the mycelium was transferred to fresh AMM containing different compounds (0.25 µg/ml voriconazol, 1 µg/ml rapamycin, 5 µg/ml monazomycin, 1 µg/ml caspofungin, 1 µg/ml trichostatin A, 0.25 µg/ml amphotericin B, 5 µg/ml AzF) or was co-cultured with *S. rapamycinicus* as described above. Mycelium was harvested using Miracloth after 1 h of incubation. The harvested mycelium was disrupted in liquid nitrogen with a precooled mortar and pestle. Ground mycelium was resuspended in phosphate buffered saline (PBS), sonicated at room temperature for 15 min and centrifuged with 15,000 rpm for 10 min. Luminescence was measured utilizing the Nano-Glo® Luciferase Assay System (Promega, Walldorf, Germany) following the manufactureŕs instruction by an infinite M200 pro microplate reader (Tecan, Männedorf, Switzerland). Luminescence of three biological replicates was measured in three technical replicates each and was normalized to the protein concentration of the supernatants. The fold change of the treatments was calculated to an untreated control.

### Identification and phylogenetic analysis of SmpR (Afu1g15910) orthologs

Orthologs of SmpR (Afu1g15910) were identified using OrthoMCL release 6.21 (Chen *et al*., 2006). 12 orthologs were excluded from the analysis because their sequence lengths were statistical outliers (length less than 200 amino acids, whereas the median sequence length for the complete ortholog set was 559 amino acids). OrthoMCL also includes multiple genomes for some species; this was reduced to a single representative genome per species, resulting in 176 orthologous sequences from 135 fungal species. Amino acid sequences were aligned using MUSCLE v3.8 (Edgar, 2004). The resulting alignment was trimmed using clipkit v2.3.0 (Steenwyk *et al*., 2020) in smart-gap mode. A maximum-likelihood phylogeny was inferred using IQ-TREE v2.2.0.3 (Minh *et al*., 2020). The ModelFinder module of IQ-TREE was used to identify JTT+F+R7 as the best substitution model for both phylogenies based on the Akaike and the Bayesian information criterion. Bootstrapping was performed using 1,000 ultrafast bootstraps. The phylogeny was visualized using ggtree 3.11.1 (Yu *et al*., 2016).

### Statistical analysis

Data and statistical analysis were performed with the GraphPad Prism 10 software package (GraphPad Software, Inc, San Diego, CA, USA). The ordinary one-way ANOVA with Tukeýs multiple comparisons test was used for significance testing of more than two groups to compare the means of each experimental group with the means of every other experimental group. Differences between the groups were considered significant at *p* ≤ 0.05. Statistical test results were included in the figure as compact letter display with different letters indicating statistical significance, while same letters mark lack of significance. Welch’s t-test was used to compare two groups. Significance was defined as a *p* value less than 0.05. Statistical test results were included in the figure legends.

## RESULTS

### The transcriptomic response of *A. fumigatus* to azalomycin F and *S. rapamycinicus*

To analyse the response of *A. fumigatus* to *S. rapamycinicus* and its inducing metabolite azalomycin F (AzF), three biological replicates of *A. fumigatus* with AzF treatment or in an *S. rapamycinicus* co-culture in three technical replicates each after 3 h of incubation were produced and a transcriptomic analysis was performed.

The co-culture with *S. rapamycinicus* significantly changed the expression of 1441 genes, with 790 genes being more transcribed in the presence of the bacterium and 651 genes being less transcribed (log_2_fold-change <> 1). The incubation with AzF resulted in an altered expression of 2065 genes, with 994 having more abundant and 1071 less abundant transcripts as compared to the untreated *A. fumigatus* Δ*KU80* wild-type control strain. The transcriptional responses of *A. fumigatus* to *S. rapamycinicus-* and AzF exposure overlapped. The abundance of 16.9 % of genes was equally lowered during AzF treatment and *S. rapamycinicus* co-culture, while 20.7 % of genes showed increased in transcription under both conditions. AzF treatment affected the abundance of 25 % genes negatively and 18 % of genes positively in a specific way, while this was the case for the expression of 8.2 % of genes with lowered and 10.1 % of genes with increased transcripts during *S. rapamycinicus* co-culture. Interestingly, transcripts of 16 genes were less abundant when the *A. fumigatus* wild-type strain was co-cultured with *S. rapamycinicus* but showed an increased abundance when the fungus was treated with AzF. By contrast, 11 genes had a low expression in AzF-treated *A. fumigatus* but higher expression when the fungus was co-cultivated with the bacterium (Figure 1, A). A list of these differentially regulated genes is shown in Table S7.

**Figure 1.**
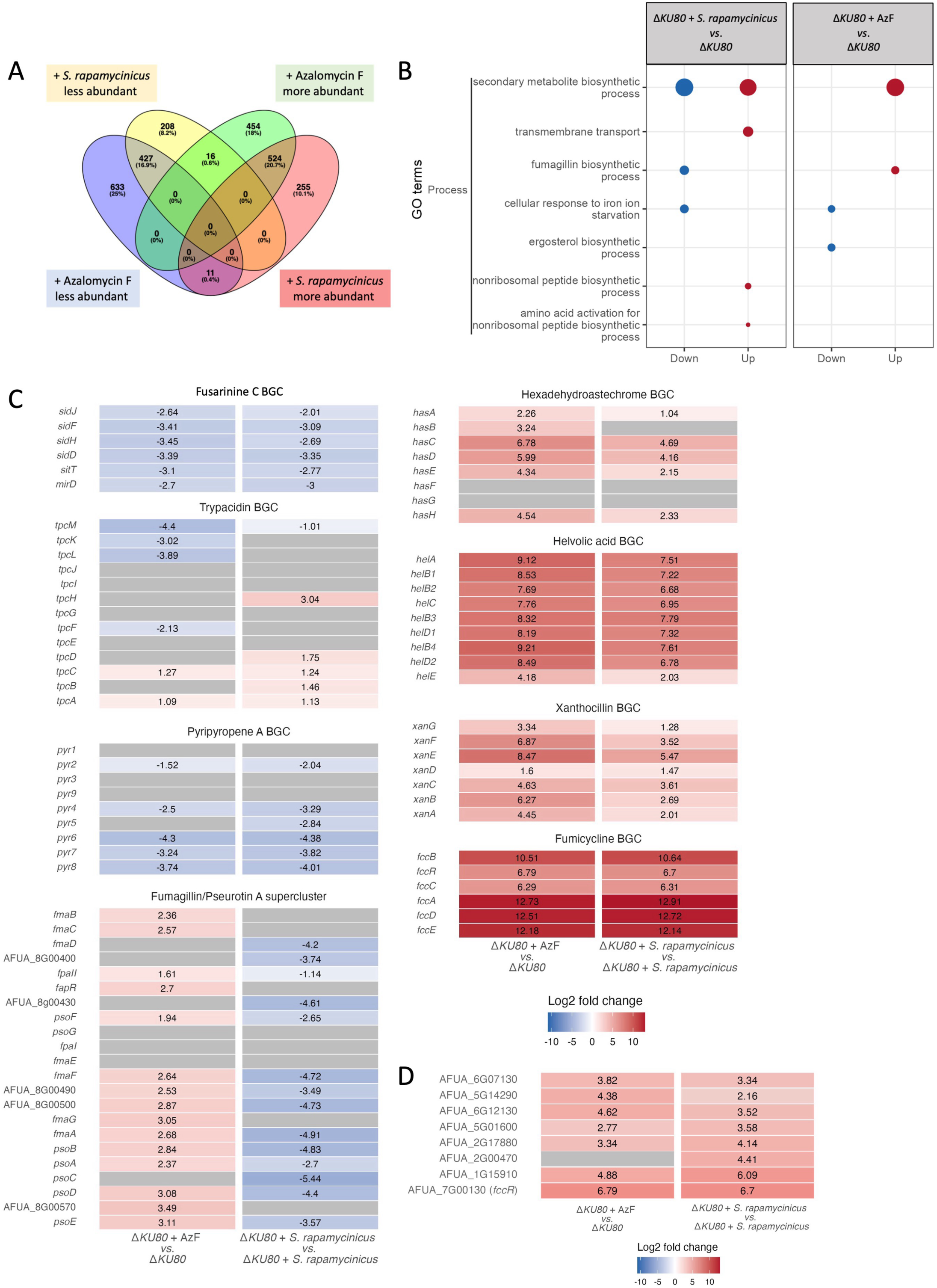
Differentially expressed genes in *A. fumigatus* co-cultured with *S. rapamycinicus* or treated with AzF. A) Venn-diagram of genes with differently abundant transcripts. *A. fumigatus* CEA17 Δ*KU80* wild type was either treated with AzF or co-cultivated with *S. rapamycinicus*. B) GO terms are depicted according to their abundance. Terms with genes which are less transcribed are indicated in blue, terms with genes which are more transcribed in red. C) Gene expression of different *A. fumigatus* BGCs. Genes belonging to a BGC are displayed with name and log_2_ fold changes during AzF treatment or *S. rapamycinicus* co-culture. The scale used defines the expression of genes from lower expressed (blue) to higher expressed (red). Genes without significant fold change are displayed in white. Grey bars indicate genes that were not found in the analysis. D) Expression of genes annotated as transcription factors. Genes with significantly increased abundance of transcripts are displayed with name and log_2_ fold changes during AzF treatment or during *S. rapamycinicus* co-culture. The scale used defines higher expression of genes in shades of red. Grey bars indicate genes that were not found in the analysis.

GO term enrichment analysis with clusterProfiler showed that most genes with altered transcription patterns belonged to the term SM biosynthetic process. Gene transcripts belonging to this term were found to be both less and more abundant (Figure 1, B). Genes with lower transcription levels under both culture conditions were more often annotated with the term cellular response to iron starvation. Additionally, genes with higher transcription were often assigned to the terms transmembrane transport, nonribosomal peptide biosynthetic process and activation of amino acids for nonribosomal peptide biosynthetic processes in the co-culture samples (Figure 1, B). KEGG term analyses also determined biosynthesis of SMs as most affected pathway, however, the term was solely annotated as less abundant in comparison to the co-culture (Figure S4). Other pathways with lower transcription belonged to the pathways of metabolism of cofactors and vitamins, lipid metabolism, carbohydrate and amino acid metabolism. Most of the KEGG terms with genes that are negatively affected were identified in *A. fumigatus* samples that had been treated with AzF (Figure S4). Pathways with higher abundant transcription were mostly related to amino acid metabolism, membrane transport and DNA replication and repair (Figure S4).

The transcription of several BGCs in *A. fumigatus* was affected under the tested conditions. The transcription of both the fusarinine C (*sid)* BGC and the pyripyropene A (*pyr)* BGC was significantly decreased in *A. fumigatus* that was treated with AzF or co-cultured with *S. rapamycinicus*. By contrast, the transcription of the hexadehydroastechrome (*has)*, the helvolic acid (*hel)*, the xanthocillin (*xan*) and fumicycline (*fcc*) BGCs were strongly increased upon AzF treatment as well as during co-culture of the fungus with *S. rapamycinicus* (**Fehler! Verweisquelle konnte nicht gefunden werden.**). The *fcc* BGC had the highest log_2_ fold changes detected for any of the highly transcribed BGCs, followed by the *hel*, *xan* and *has* BGCs. In case of the trypacidin BGC and the intertwined fumagillin/pseurotin A (*fma*) supercluster, the different culture conditions led to differential gene expression. While the treatment with AzF resulted in increased transcription of the *fma* BGC and a decreased transcription of the trypacidin, the co-culture with *S. rapamycinicus* caused a decreased transcription of the supercluster and an increased transcription of the trypacidin BGC (Figure 1, C). Functional annotations of the genes belonging to the aforementioned BGCs can be found in Table S8.

The transcriptome data suggested that both *S. rapamycinicus* and AzF trigger the activation of BGCs in *A. fumigatus* similar to that observed for the co-incubation of *A. nidulans* with *S. rapamycinicus* or the treatment of *A. nidulans* with AzF (Krespach *et al*., 2023). Transcripts of the *fcc* BGC showed the highest increase in abundance. This led us to hypothesize that potential global regulators of the *fcc* BGC are induced by both the bacterium and AzF. Therefore, we investigated the expression pattern of several transcription factor genes. Genes annotated as C6 transcription factor, putative fungal transcription factor, DNA-binding transcription factor activity that modulates the transcription of specific gene sets transcribed by RNA or with predicted DNA binding in FungiDB (Alvarez-Jarreta *et al*., 2023), were grouped based on their log_2_ fold changes. Only genes with increased transcription were analysed, since anticipated regulators of the *fcc* BGC should have an increased expression under AzF treatment and during *S. rapamycinicus* co-cultivation. The top five highest upregulated transcription factor genes of each treatment were analysed. The transcription factor with the highest increase of transcription in both conditions was *fccR*, the *fcc* cluster-encoded regulator. The gene AFUA_1g15910 showed fold changes in transcription levels comparable to the fold changes of *fccR* under both treatments (Figure 1, D). This gene was previously described in the context of mitochondrial dynamics and was named *mdd2.* Furthermore, in an overexpression study of zinc cluster transcription factors in *A. fumigatus*, it was designated as *zcf63* (Sturm *et al*., 2020; Sasse *et al*., 2023). However, given its possible involvement in regulation of several BGC, we renamed the gene as <u>s</u>econdary metabolite <u>m</u>ultiple <u>p</u>athway regulator (*smpR*).

### SmpR is a fungal zinc cluster transcription factor induced by distinct molecular triggers

To characterize the structural properties of SmpR, its structure was modelled with the AlphaFold3 software (Abramson *et al*., 2024). SmpR belongs to the group of zinc cluster transcription factors, a group of proteins commonly and only found in fungi which contain a unique Zn(II)_2_Cys_6_ motif (MacPherson *et al*., 2006). Common structures of these proteins are an N-terminal DNA binding domain (IPRO001138), and two alpha helices harbouring two zinc atoms bound by six cysteine residues (Gardner *et al*., 1991; Pan and Coleman, 1990). The DNA binding domain for SmpR spans amino acid 20 to 47 which is followed C-terminally by a dimerization domain. The following variable central region is responsible for transcriptional activation (Poch, 1997, MacPherson *et al*., 2006). In addition to all of these conserved features that are specific to fungal zinc cluster transcription factors, InterPro analysis (Blum *et al*., 2024) also revealed the presence of a conserved, so called “fungal_trans” domain in the central region of SmpR, which can act in the activation of zinc cluster transcription factors (Mayer *et al*., 2023; Poch, 1997) (Figure 2, A).

**Figure 2.**
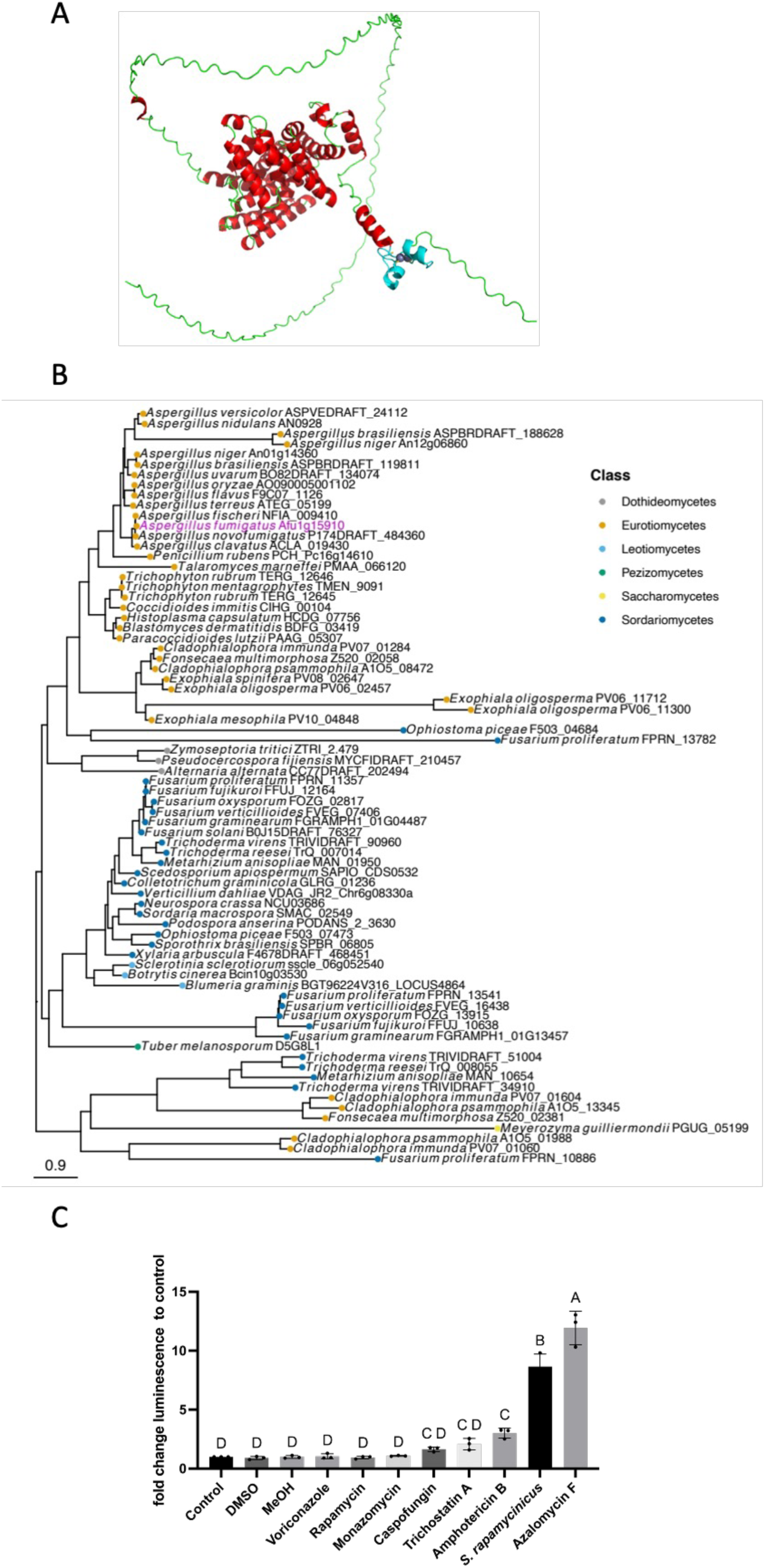
*SmpR* encodes a specifically induced fungal zinc cluster transcription factor. A) AlphaFold3 model of SmpR. The zinc cluster motif is located from amino acid 20 to 47 (in cyan). B) Maximum likelihood phylogeny of selected *A. fumigatus smpR* (Afu1g15910) orthologs. The *A. fumigatus* ortholog is indicated in purple text. The colour at the end of the branches denotes the fungal class to which the organism belongs. C) Luminescence fold change in cultures of the *smpR-nluc* strain incubated with different compounds compared to the untreated control.

Phylogenetic analysis revealed that *smpR* orthologues are limited to the phylum Ascomycota and the classes Dothideomycetes, Eurotiomycetes, Leotiomycetes, Orbiliomycetes, Pezizomycetes, Saccharomycetes and Sordariomycetes. The class of Eurotiomycetes harboured most of the *smpR* orthologues, followed by the class of Sordariomycetes, while only two orthologues of *smpR* could be found in the class of Saccharomycetes. Most genomes encoded a single copy of *smpR*, but several phyto-pathogenic *Fusarium* spp encoded two copies, suggesting a beneficial role and/or neofunctionalization. As expected, the most closely related orthologs of *smpR* (syn: Afu1g15910) were found in the family *Aspergillaceae*, in both human pathogenic- and non-pathogenic species (Figure 2, B).

To determine the conditions which lead to activation of *smpR* expression, a nanoluciferase tag was fused to the C-terminus of SmpR and the corresponding recombinant *A. fumigatus* strain expressing this fusion was incubated in the presence of different compounds or co-cultivated with different bacteria. Only in co-culture with *S. rapamycinicus*, a producer of AzF, or the addition of 5 µg/ml pure AzF the expression of the *smpR* gene fusion was significantly increased (Figure 2, C). Notably, the addition of pure AzF led to an even stronger increase of *smpR* expression than observed during the fungal-bacterial co-culture. Of the other tested SMs from Streptomycetes like the macrolide rapamycin, the cyclic arginoketide monazomycin, the hydroxamic acid trichostatin A and the polyene macrolide amphotericin B, only amphotericin B was able to induce the expression of *smpR* in comparison to an DMSO control culture. Amphotericin B is also known to induce fumicycline production in *A. fumigatus* (Abou-Kandil *et al*., 2024) although to a lesser extent than observed during co-cultures with *S. rapamycinicus.* This indicates that the induction of *smpR* expression is restricted to specific molecular triggers.

### *smpR* is involved in regulating fumicycline production

For a more detailed investigation of the function of SmpR we deleted the corresponding gene in *A. fumigatus* Δ*KU80* by homologous recombination. To assess the phenotypic changes of the deletion mutant Δ*smpR*, the growth of the wild-type Δ*KU80* strain and the Δ*smpR* mutant strain on different media was determined. There was no significant difference between the colony diameters during growth on minimal (AMM) agar or rich medium (Malt agar) (Figure S5, A). There were no observable differences in growth between the Δ*smpR* mutant strain and the Δ*KU80* wild type on AMM agar plates containing compounds inducing cell wall- (30 µg/ml congo red, 30 µg/ml calcofluor white [CFW] and 0.5 µg/ml caspofungin), membrane-disruption (0.5 µg/ml voriconazole and 0.5 µg/ml amphotericin B), osmotic- (1 M NaCl) and oxidative stress-inducing compounds (4 mM H_2_O_2_ and 4 µM menadione) or AzF-containing agar plates (10 µg/ml) (Figure S5, B). To test the sensitivity of the deletion strain Δ*smpR* against AzF and the antimycotic agents amphotericin B, voriconazole and caspofungin their minimal inhibitory concentration (MIC) was determined according to the broth dilution methodology for conidia-forming moulds (EUCAST, 2020). There was no increase in susceptibility or resistance against either of the compounds tested (Table S 9). However, when co-cultured with *S. rapamycinicus,* Δ*smpR* was not able to produce significant levels of the intense yellow colour as the Δ*KU80* wild type. This colour change is indicative of the production of fumicyclines, observed when the fungus is co-cultured with this bacterium (Figure 3, A) (König *et al*., 2013). In accordance, the *smpR* mutant strain produced less fumicyclines A, B and C (Figure 3, C *top*). To determine the effect of *smpR* deletion on the expression of the fumicycline BGC, qRT-PCR was performed. This analysis showed that all *fcc* BGC genes were significantly less transcribed in Δ*smpR* (Figure 3, B *top*). The effect of decreased abundance of *fcc* transcripts was restored when the *smpR* deletion strain was complemented with the *smpR* gene. All *fcc* genes, had transcript levels comparable to the level determined in co-culture of the Δ*KU80* wild type with *S. rapamycinicus* (Figure 3, B *top*). The complementation strain was able to produce Δ*KU80-*like levels of fumicycline A, B and C that could also be detected visibly and by LC-MS analysis (Figure 3, A and C *top*). To test the biological effects of an *smpR* overexpression, we generated an *A. fumigatus* strain expressing *smpR* under control of the *xylP* promoter, *Xyl*_P_-*smpR*. When induced by substitution of glucose by xylose in AMM, *Xyl*_P_-*smpR* produced fumicyclines independently of the co-cultivation with *S. rapamycinicus* (Figure 3, A and C *bottom*). However, the xylose-induced overexpression of *smpR* only triggered a lower production of fumicyclines A, B and C as compared to Δ*KU80* wild type in co-culture with *S. rapamycinicus* (Figure 3, C *bottom*). The addition of *S. rapamycinicus* to the induced *smpR*-overexpressing strain still caused a significantly higher expression of all *fcc* cluster genes (Figure 3, B *bottom*) but only a slightly increased production of all fumicycline isomers (Figure 3, C *bottom*). The decrease in fumicycline production which was accompanied by a reduced expression of *fcc* cluster genes in the Δ*smpR* deletion strain and the slightly increased fumicycline production in *Xyl*_P_-*smpR* indicated that *smpR* is important for the regulation of the fumicycline cluster, but does not represent the only regulatory element required for fumicycline production.

**Figure 3.**
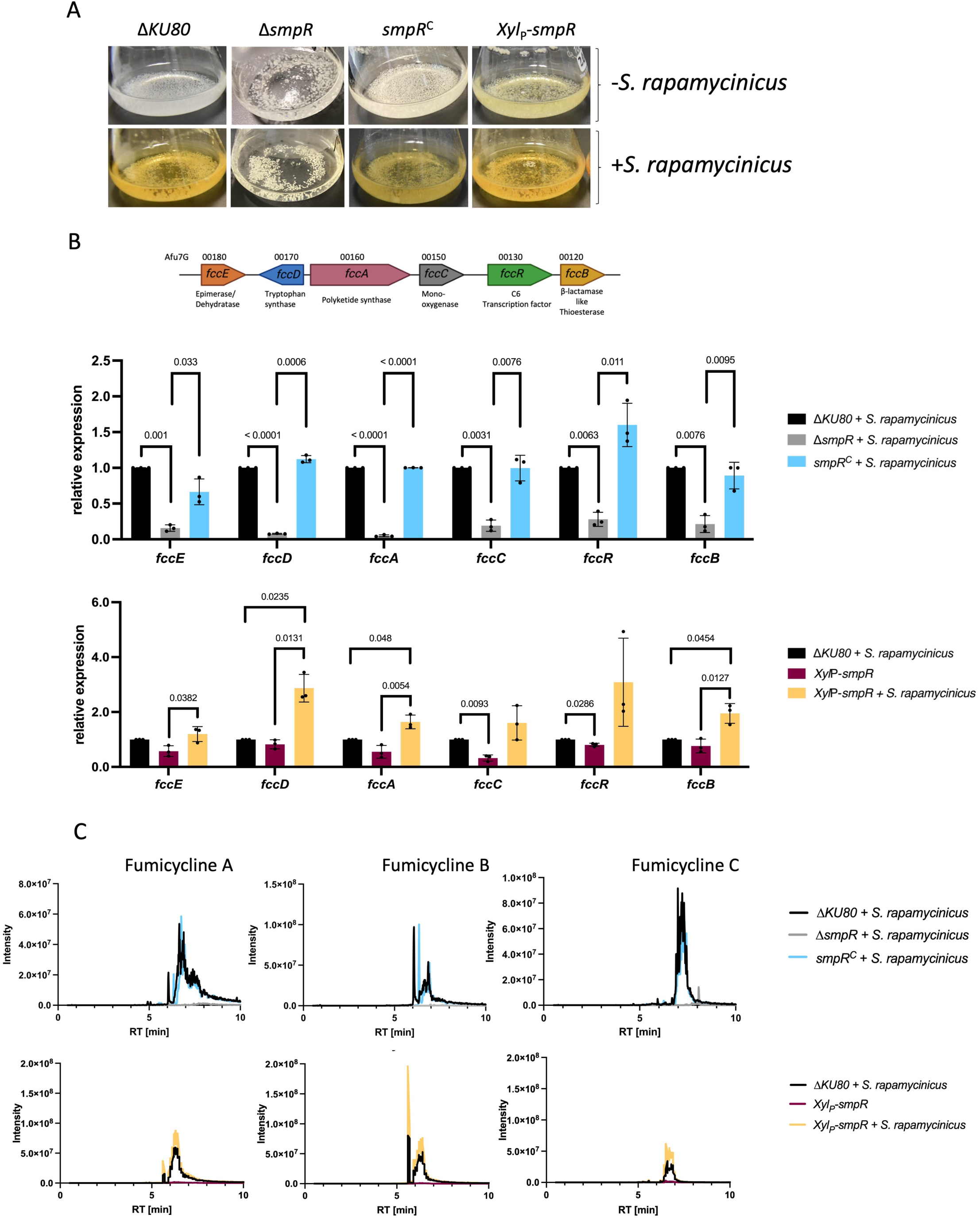
Analysis of *fcc* BGC gene expression and fumicycline production. A) Co-cultures with *S. rapamycinicus* or axenic cultures of Δ*KU80*, Δ*smpR, smpR*^C^ and *Xyl*_P_-*smpR*. *Xyl*_P_-*smpR* was induced by the addition of 1 % (w/v) xylose to the culture medium. B) *top* Schematic representation of the *fcc* cluster genes with annotations of putative activities. *Down* Determination of the relative expression of *fcc* genes of these strains by qRT-PCR. Expression of target genes was normalised to *act1* expression using the 2^-ΔΔCt^ method. The expression level of target genes was compared to the expression of the target genes in the Δ*KU80* wild type which was co-cultured with *S. rapamycinicus*. C) Extracted ion chromatograms of Δ*KU80*, Δ*smpR, smpR*^C^ during co-culture with *S. rapamycinicus,* xylose-induced *Xyl*_P_-*smpR* and xylose-induced *Xyl*_P_-*smpR* in co-culture with *S. rapamycinicus*.

### *smpR* acts hierarchically upstream of *fccR*

Previously, the overexpression of *fccR* has been shown to induce and upregulate the expression of the *fcc* cluster genes as well as the production of fumicyclines independently of a co-cultivation (König *et al*., 2013). In agreement, *fcc* cluster gene expression and fumicycline production in the Δ*fccR* mutant strain were abolished, indicating the important role of *fccR* for activation of the *fcc* cluster (König *et al*., 2013; Chooi *et al*., 2013; König, 2015). However, the factors leading to activation of *fccR* expression remained obscure. Therefore, we investigated the relationship between SmpR and FccR. For this purpose, we overexpressed *smpR* in the Δ*fccR* mutant strain as well as *fccR* in the Δ*smpR* mutant strain. While the overexpression of *smpR* in Δ*fccR* did not lead to formation of the characteristic yellow colouring of the culture medium, indicating fumicycline production, the overexpression of *fccR* in Δ*smpR* did, even surpassing the amounts reached in the Δ*KU80* wild type observed in co-culture with *S. rapamycinicus*. The addition of *S. rapamycinicus* did not enhance fumicycline production in either of the strains (Figure 4, A and C). Expression of *fccA,* analysed as a representative gene of the *fcc* BGC, was abolished in the *Xyl*_P_-*smpR* Δ*fccR* strain while its expression was significantly higher in the *Xyl*_P_-*fccR* Δ*smpR* strain as compared to the Δ*KU80* wild type co-cultured with *S. rapamycinicus* (Figure 4, B). While an overexpression of *smpR* can induce *fccR* expression independently of *S. rapamycinicus* co-culture and the deletion of *smpR* led to a significant decrease in expression of *fccR*, an overexpression of *smpR* cannot overcome an *fccR* deletion on fumicycline production. This suggests that *smpR* is involved in the activation of *fccR* expression upstream of the *fcc* cluster.

**Figure 4.**
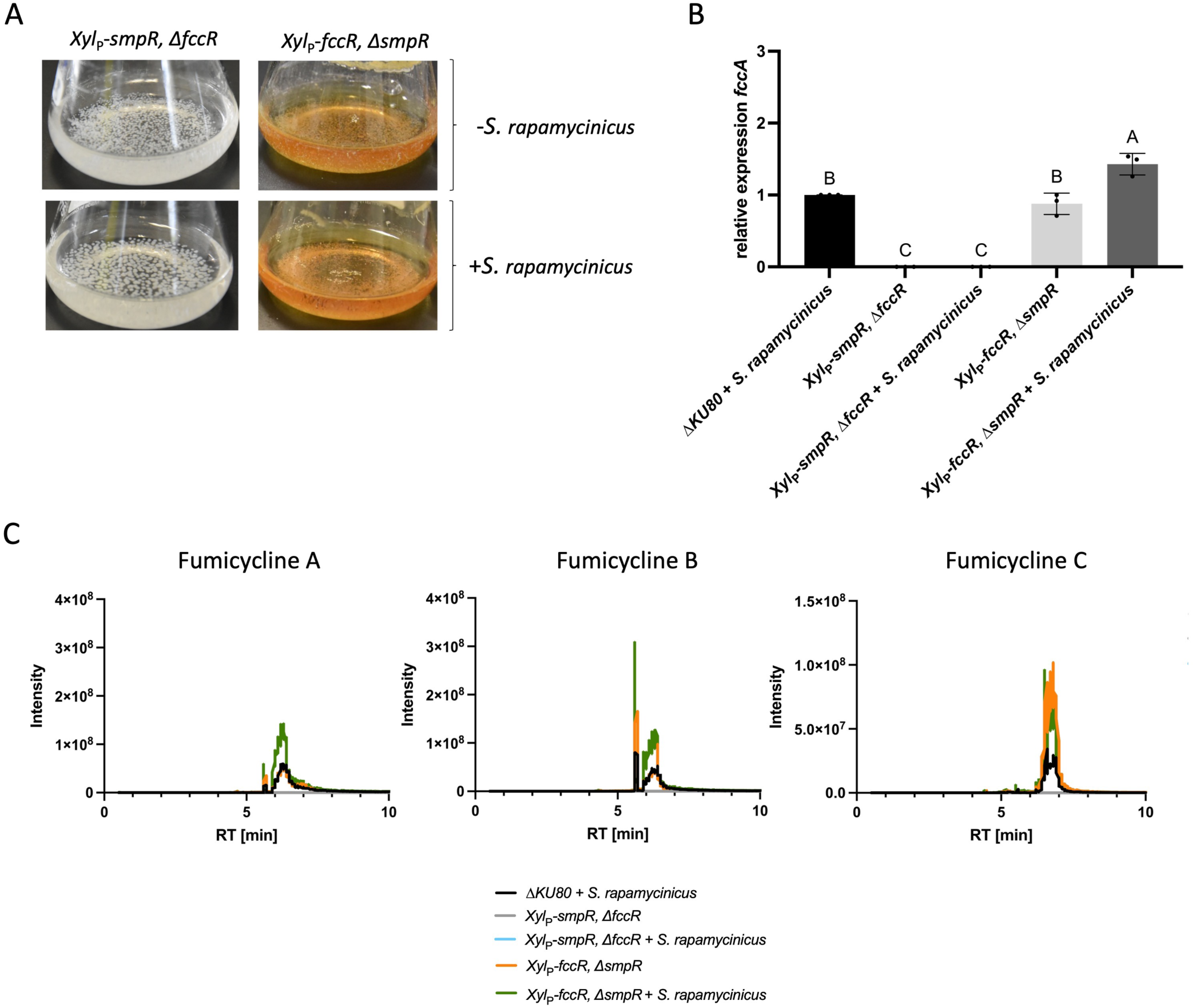
Relationship between *smpR* and *fccR.* A) Co-cultures with *S. rapamycinicus* and axenic cultures of *Xyl*_P_-*smpR*, Δ*fccR* and *Xyl*_P_-*fccR,* Δ*smpR*. Both overexpressing strains were induced by the addition of 1 % (w/v) xylose to the culture medium. B) Determination of the relative expression of *fccA* by qRT-PCR. Expression of target genes was normalised to *act1* expression using the 2^-ΔΔCt^ method. The expression level of target genes was compared to the expression of the target genes in Δ*KU80* which was co-cultured with *S. rapamycinicus*. C) EICs of *Xyl*_P_-*smpR*, Δ*fccR* and *Xyl*_P_-*fccR,* Δ*smpR* in axenic culture in xylose-AMM as well as in xylose-AMM + *S. rapamycincus*.

### *SmpR* deletion affects expression of genes associated with secondary metabolism and an antibacterial response

To analyse the effect of the deletion of *smpR* on the transcriptome of *A. fumigatus* treated with AzF or co-cultivated with *S. rapamycinicus*, transcriptomic analysis was performed. Such an analysis may also reveal regulatory targets of *smpR* besides the *fcc* cluster. The deletion of *smpR* led to a decreased transcription of 173 genes in AzF-treated samples and to a reduced transcription of 400 genes in co-cultures with *S. rapamycinicus*. 17 genes were more abundant in Δ*smpR* under AzF treatment and 136 genes showed higher transcript levels when Δ*smpR* was co-cultured with *S. rapamycinicus*. GO term enrichment analysis of gene sets that showed an altered abundance in Δ*smpR*, irrespective of treatment, revealed an enrichment of genes involved in secondary metabolic process and transmembrane transport. Additional KEGG terms showed a decrease of transcription of genes belonging to the term metabolism of tyrosine and phenylalanine (Figure 5, A, B). Increased transcript levels were found in the terms of IgE binding, phosphopantetheine binding in AzF-treated samples. In *S. rapamycinicus* co-cultures, increased levels of transcripts were found associated with heme binding, fungal-type cell wall as well as intracellular anatomical structure (Figure 5, A).

**Figure 5.**
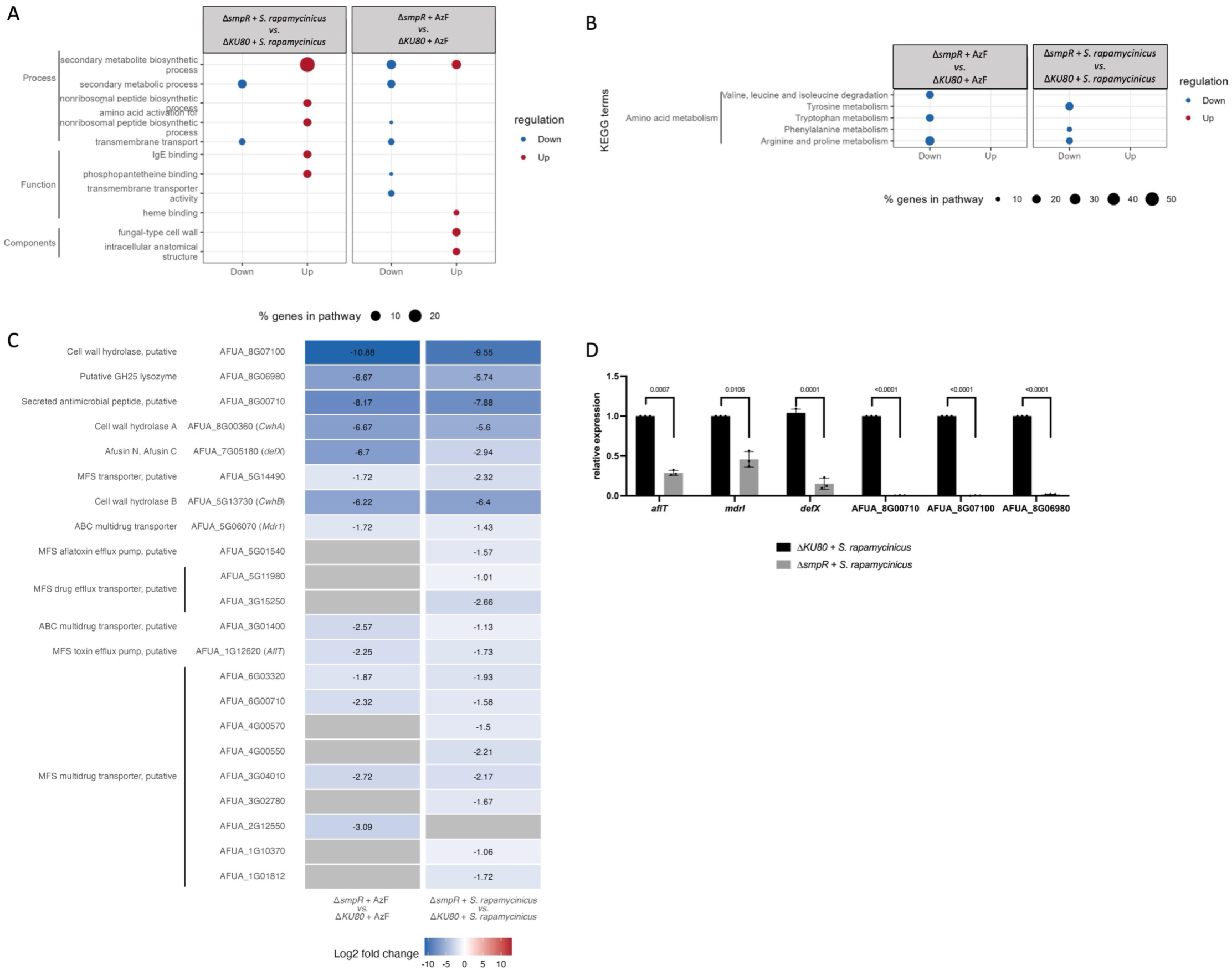
Gene expression in Δ*smpR deletion strains.* A) GO terms are depicted according to their abundance. Lower transcribed terms are labelled in blue, higher transcribed terms in red. B) KEGG terms show lowered transcriptions of terms in blue. C) Genes with significantly lowered transcription in Δ*smpR* are displayed with name, annotation and log_2_ fold changes under AzF treatment or during *S. rapamycinicus* co-culture. The scale used defines lower expression of genes in shades of blue. Genes without significant fold change are displayed in white. Grey bars indicate genes that were not found in the analysis. D) Expression analysis of selected genes potentially associated with antibacterial defence. Relative expression of *aflT, mdrI, defX*, AFUA_8G00710, AFUA_8G07100 and AFUA_8G06980 by qRT-PCR. Expression of target genes in mutant strain Δ*smpR* was normalised to *act1* expression using the 2^-ΔΔCt^ method. The expression level of target genes was compared to the expression of the target genes in the Δ*KU80* wild type co-cultured with *S. rapamycinicus*.

Since the loss of *smpR* had a substantial effect on fumicycline production and fumicyclines exhibit antibacterial activity, we investigated the antibacterial activity in more detail. Therefore, we searched for gene annotations connected to antibacterial defence and compared the response of Δ*smpR* and Δ*KU80* in co-culture and under AzF treatment. The transcription of genes annotated as drug transporters (MFS and ABC-type), antimicrobial peptides as well as bacterial cell wall hydrolases and lysozyme were significantly decreased in Δ*smpR* (Korczynska *et al*., 2010; Machata *et al*., 2024). The response to AzF and *S. rapamycinicus* mostly overlapped (Figure 5, C). Genes encoding for MFS toxin efflux pump AflT, the ABC transporter Mdr1, as well as *defX* and AFUA_8G00710 which encode antibacterial peptides, as well as genes encoding the putative cell wall hydrolase AFUA_8G07100 and the putative lysozyme AFUA_8G06980 were further analyzed by qRT-PCR (Korczynska *et al*., 2010; Machata *et al*., 2024). While transcription of all selected genes was induced during co-culture (data not shown), qRT-PCR analysis showed that transcript levels of the drug transporters AflT and Mdr1 as well as *defX* were significantly lower in the Δ*smpR* mutant strain. Deletion of *smpR* led to an almost completely absence of transcripts of genes AFUA_8G00710, AFUA_8G07100 and AFUA_8G06980 (Figure 5, D).

### *SmpR* deletion affects the transcription of multiple BGCs

The deletion of *smpR* did not only affect the transcription of the fumicycline BGC but also of parts of the *has* BGC and all genes in the *hel* and *xan* BGCs (Figure S6, A). The BGCs are known to encode genes responsible for the production of antimicrobial SMs (Raffa *et al*., 2021; König *et al*., 2013, Sang *et al*., 2024). The lowered abundance of transcription of the *hel* and *xan* BGC was observed in the Δ*smpR* mutant strain that was treated with AzF as well as in co-culture with *S. rapamycinicus* (Figure S6, A middle and bottom). The lowered abundance of helvolic acid production was in part confirmed by LC-MS analysis revealing that the level of helvolic acid in the co-culture supernatant of Δ*smpR* and *S. rapamycinicus* was significantly lower as compared to co-cultures with the Δ*KU80* wild-type strain (Figure S6, B). The effect on the *has* BGC was only observed in Δ*smpR* samples that were co-cultured with *S. rapamycinicus* but not when AzF was added (Figure S6, A top). To further investigate the effect of SmpR on these BGCs, qRT-PCR analysis of the *smpR* deletion mutant and the *smpR*-overexpression strain was conducted. The deletion of *smpR* led to a significantly reduced transcription of all genes of the *hel* and *xan* cluster as well as a reduced expression of *hasH, hasE, hasD* and *hasC* (Figure 6, A, B and C *top*). By contrast, overexpression of *smpR* increased transcription of all aforementioned genes, especially of the *xan* cluster, where transcription levels increased 50- to 100-fold for some genes of the clusters. For the *has* and *hel* BGCs a more moderate increase was observed (Figure 6, A, B and C *bottom*). The addition of *S. rapamycinicus* to the smpR-overexpressing strain of *A. fumigatus* increased the expression of *hasH, hasE, hasD* and *hasC* even further up to 40-fold in comparison to the mono-culture of the induced *smpR* overexpressing strain. This effect was not observed for the *xan* and *hel* BGC, where addition of *S. rapamycinicus* only led to a modest increase in expression of the cluster genes in comparison to the mono-culture of the *smpR* overexpression strain or even a significantly lowered transcription (Figure 6, A, B and C *bottom*). Together. this finding indicates that SmpR also acts as more global regulatory element at least also for the *has, hel* and *xan* BGCs.

**Figure 6.**
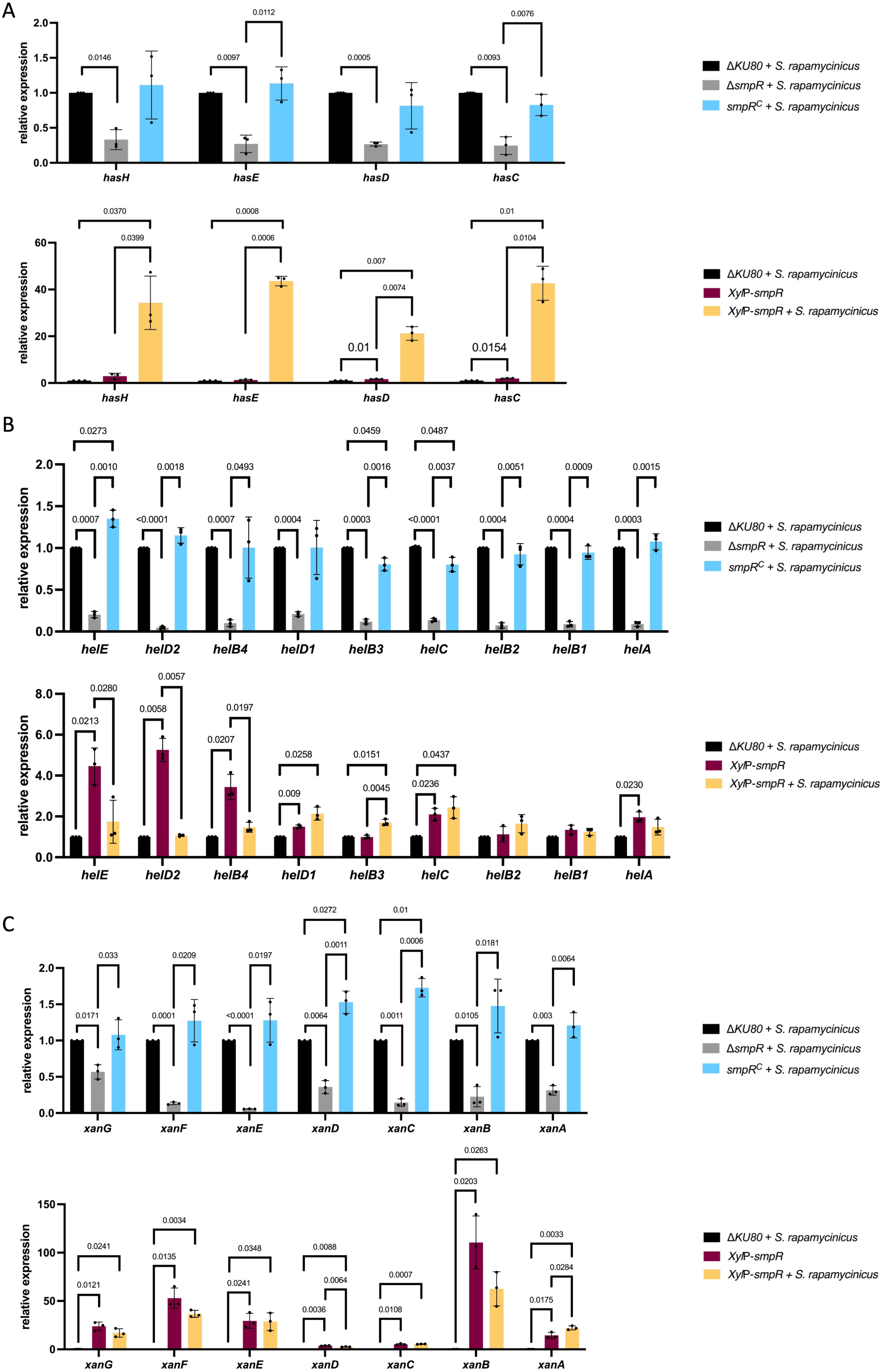
Transcript levels of the *has*, *hel* and *xan* BGC genes in *smpR* mutant strains. A) Transcript levels of the *has* BGC genes in Δ*smpR* and *Xyl*_P_-*smpR* mutant strains. B) Transcript levels of the *hel* BGC genes in Δ*smpR* and *Xyl*_P_-*smpR* mutant strains C) Transcript levels of the *xan* BGC genes in Δ*smpR* and *Xyl*_P_-*smpR* mutant strains. Transcript levels of target genes in mutant strain Δ*smpR* was normalised to *act1* expression using the 2^-ΔΔCt^ method. The expression level of target genes was compared to the expression of the target genes in the Δ*KU80* wild type co-cultured with *S. rapamycinicus*.

### SmpR-regulated fumicycline production led to inhibition of bacterial growth

To investigate whether soil-dwelling bacteria other than *S. rapamycinicus*, can induce the transcription of *smpR* and the SmpR-regulated BGCs, we isolated different bacteria from local soil samples. The isolation yielded *Arthrobacter* spp. and *Kribbella* spp. which we choose specifically for the analysis since both species are known to produce SMs with antifungal properties (Ramlawi *et al*., 2021; Virués-Segovia *et al*., 2022). We further examined the potential of the SmpR regulated BGCs to exert antimicrobial activity against these isolates. The gram-positive and gram-negative laboratory strains *B. subtilis* 168 and *E. coli* DH5α, respectively, were used as controls. The co-cultures of the bacteria with *A. fumigatus* were extracted for LC/MS analysis as well as harvested for qRT-PCR. All tested bacterial isolates and strains were able to induce the transcription of *smpR*, although to a lesser extent than *S. rapamycinicus* (Figure 7, A; Figure S7). The soil isolates *Kribbella* spp. and *Arthrobacter* spp. were also able to induce the transcription of the fumicycline BGC genes *fccA* and *fccR*, whose transcription levels were analysed as representatives of the fumicycline BGC as well as fumicycline production (Figure 7, A, B). None of the tested co-cultures of *A. fumigatus* and soil isolates or laboratory strains showed expression of representative genes of the *has*, *hel* or *xan* BGCs (data not shown).

**Figure 7.**
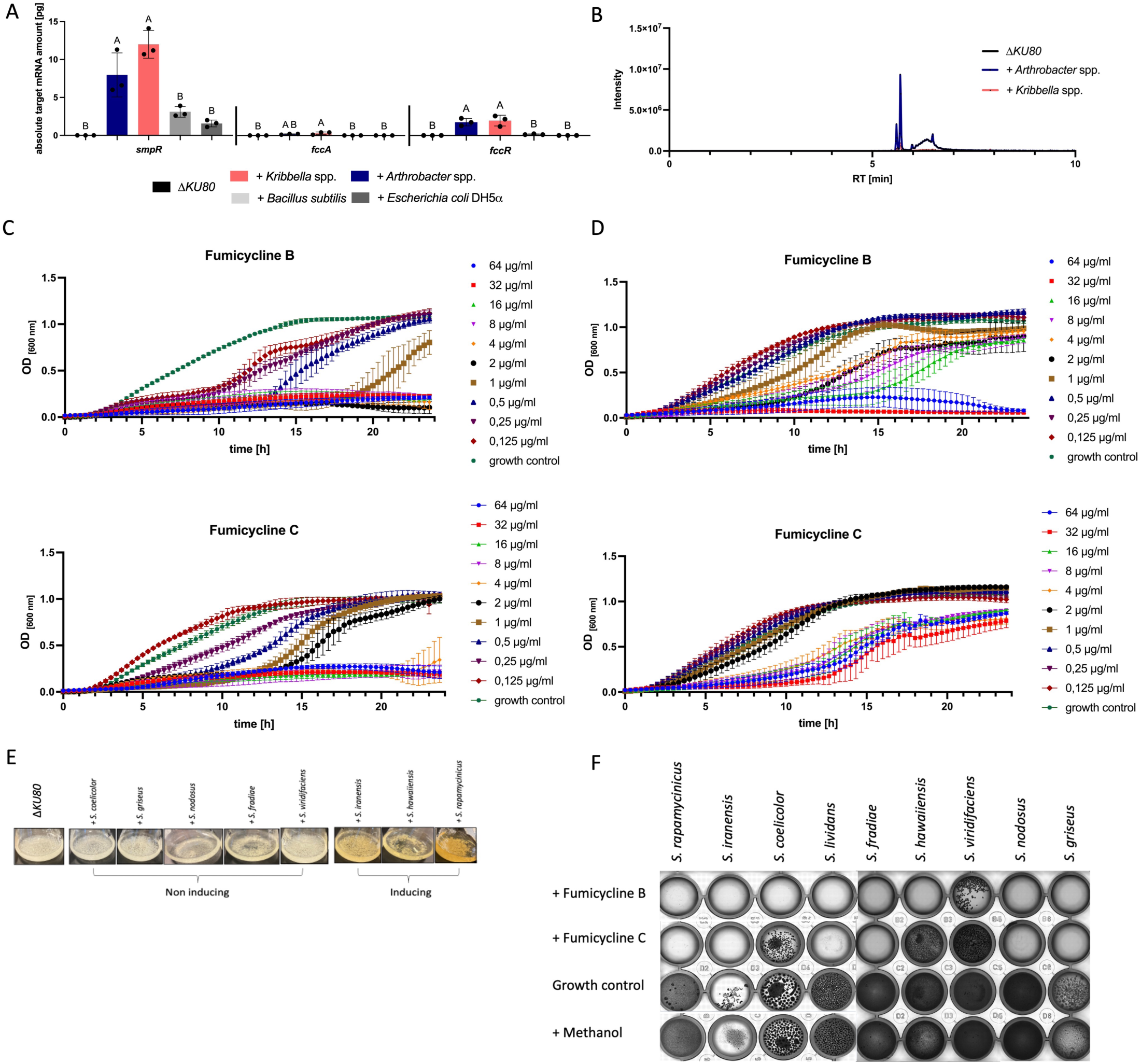
Induction of production of fumicyclines and their effect on bacterial growth. A) Analysis of transcript levels of different genes during co-cultures of *A. fumigatus* with different soil isolated bacteria and laboratory strains. The transcript level of *smpR, fccA* and *fccR* was determined *via* qRT-PCR as absolute mRNA quantification. B) EICs of fumicycline B in Δ*KU80* in axenic culture and as co-cultures with *Arthrobacter* spp. and *Kribbella* spp. C) Growth curves of *Arthrobacter* spp. with different concentration of *top* fumicycline B and *bottom* fumicycline C. D) Growth curves of *Kribbella* spp. with different concentration of *top* fumicycline B and *bottom* fumicycline C. E) Co-cultures of *S. coelicolor, S. griseus, S. nodosus, S. fradiae, S. viridifaciens, S. iranensis, S. hawaiiensis* and *S. rapamycinicus* with *A. fumigatus* Δ*KU80*. F) The indicated *Streptomyces* species were grown in the presence of 64 µg/ml fumicycline B or fumicycline C in 48-well plates. Growth was measured microscopically after 24 h for 96 h. Untreated wells and wells containing methanol served as controls.

Recently, it was reported that neosartoricins exhibit an antiproliferative activity on CD3/CD28-activated murine spleenic T-cells and no inhibitory effect on the tested Gram-positive or Gram-negative bacteria (Chooi *et al*., 2013). Since the induction of *smpR* and the production of fumicyclines (neosartoricins) are directly linked to the co-culture with different bacteria, an inhibitory effect on the inducing bacteria seemed likely. In agreement, fumicyclines showed moderate growth inhibitory activity against *S. rapamycinicus* in agar diffusion tests (König *et al*., 2013). To further evaluate the activity of fumicyclines on the *fcc* BGC-inducing *Kribbella spp. and Arthrobacter spp.* and to compare the effect of different fumicycline isomers on these bacteria, we measured bacterial growth in the presence of different concentrations of fumicycline B and fumicycline C. Particularly for *Arthrobacter* spp. a pronounced effect of both fumicycline isomers was detected. The bacterium exhibited no growth at fumicycline B concentrations exceeding 1 µg/ml and at lower concentrations growth delay was observed. For fumicycline C, delayed growth was visible with concentrations under 2 µg/ml and growth arrest for concentrations of 4 µg/ml and above (Figure 7, B). For *Kribbella* spp., the inhibitory effect of the fumicyclines was less pronounced. Only high concentrations (64 µg/ml and 32 µg/ml) of fumicycline B completely arrested the growth of the bacterium, while lower concentrations between 16 µg/ml and 1 µg/ml led to a delayed onset of the exponential growth phase. For fumicycline C, all concentrations above 2 µg/ml induced slower growth as compared to concentrations under 2 µg/ml and the growth control (Figure 7, C).

Since the growth of *S. rapamycincius* is not reliably traceable with the measurement of optical density, we cultivated spores of the bacterium in the presence of 64 µg/ml pure fumicycline B or fumicycline C for 96 h in 48-well plates. We also included other *Streptomyces* species to analyse their ability to induce the production of fumicyclines. Therefore, we co-cultured *Streptomyces* species of different clades, *i.e*., *S. coelicolor, S. griseus, S. nodosus, S. fradiae, S. viridifaciens, S. hawaiiensis* as well as the known inducers *S. iranensis* and *S. rapamycinicus* with *A. fumigatus* Δ*KU80* wild type and compared their ability to induce the production of fumicyclines by visible observation. The majority of the strains tested here were not able to induce the production of fumicyclines, visible by the lack of the yellow colouring of the culture supernatant (Figure 7, D). Only *S. hawaiiensis* and the known inducers of fumicycline production, *S. iranensis* and *S. rapamycinicus*, which are phylogenetically distant to *S. hawaiiensis*, induced the production of fumicyclines (Figure 7, D). However, the ability of almost all of the tested spores of *Streptomyces* species to produce mycelia was inhibited by fumicyclines as compared to the methanol growth control (Figure 7, E). Fumicycline B had a stronger inhibitory effect than fumicycline C. Only *S. viridifaciens* was able to grow after 96 h of incubation in the presence of fumicycline B, while *S. coelicolor, S. hawaiiensis* and *S. viridifaciens* were able to grow in the presence of fumicycline C after 72 h (Figure 7, E). This effect of growth inhibition was reversible and not only observable with spores of the tested species but also with their mycelium (data not shown). Together, fumicyclines are potent inhibitors of growth of different soil-dwelling bacterial species. However, the strength of the inhibitory effect differs between the bacterial species.

## DISCUSSION

Environmental signals can alter the transcriptome of fungi and as a consequence, they can adapt and respond to environmental cues. These signals can be represented by low molecular weight compounds including SMs that are produced and exchanged by microorganisms. These compounds can have profound effects on neighbouring microorganisms including the induction of the expression of silent BGCs (Netzker *et al*., 2018). Here, we defined a specific response of *A. fumigatus* to the treatment of the fungus with AzF as well as to its co-cultivation with *S. rapamycinicus.* Several BGCs were either induced or repressed by the two stimuli, demonstrating that AzF also acts as a stimulus for SM production in *A. fumigatus* as reported for *A. nidulans* (Krespach *et al*., 2023). Although *A. fumigatus* is found in the environment, the fungus is also able to infect humans by overcoming the host defence e.g. through production of SMs that modulate the host immune response (Scharf *et al*., 2014). Several BGCs that are likely related to virulence and host adaptation in *A. fumigatus*, like the extracellular siderophore fusarinine C and the inhibitor of the acyl-CoA:cholesterol acyltransferase pyripyropene A showed reduced transcription levels under AzF treatment and in an *S. rapamycinicus* co-culture (Omura *et al*., 1993; Schrettl *et al*., 2007; Itoh *et al*., 2010; Bignell *et al*., 2016). The transcription level of the BGC-encoding genes for production of the immunomodulatory compounds fumagillin and pseurotin A, was lower when *A. fumigatus* was co-cultured with *S. rapamycinicus* and stronger during AzF treatment (Ishikawa *et al*., 2009; Fallon *et al*., 2010; Wiemann *et al*., 2013). A similar effect was seen for the antiphagocytic trypacidin BGC (Mattern *et al*., 2015). While transcription of several genes of the BGC was increased in the presence of *S. rapamycinius,* treatment with AzF lowered the expression of these genes. This suggests that *S. rapamycinicus* has additional factors that have an influence on differential BGC expression, as compared to the pure metabolite AzF.

Fungal-specific zinc cluster transcription factors are key elements in regulating drug tolerance, adaptive processes to different stressors and activation of defence mechanisms against competitors as well as in regulation of SMs in *A. fumigatus* (Brakhage, 2013; Hagiwara *et al*., 2017; Valero *et al*., 2021; Ries et al., 2020). Here, we identified the transcription factor-encoding gene *smpR* as a regulator of multiple BGCs encoding antibacterial SMs and further antibacterial defence-related genes in *A. fumigatus*. For the latter, we found that the expression of genes encoding multidrug transporters like Mdr1 and AflT, and of antibacterial peptides were significantly less transcribed in the Δ*smpR* deletion mutant. Furthermore, the expression of genes encoding cell wall hydrolases and lysozyme was fully abolished when SmpR was absent, which suggests that these genes are targets of SmpR (Korczynska *et al*., 2010, Machata *et al*., 2024). Remarkably, the deletion of *smpR* not only influenced the transcript levels of single genes that are potentially associated with the antimicrobial response of the fungus but also reduced the transcript levels of some genes of the *has* BGC, and of all genes of the *hel, xan* and *fcc* BGCs. In agreement, the production of all fumicycline isotypes as well as helvolic acid was decreased in the Δ*smpR* mutant. The loss of SM production after deletion of the transcription factor gene *smpR*, that is not located within a BGC, has already been reported. An example is the transcription factor RglT, which regulates the gliotoxin biosynthesis genes in *A. fumigatus* but is not part of the *gli* BGC. The loss of RglT resulted in the lack of gliotoxin production and thereby had a similar effect as deletion of the cluster-encoded transcription factor *gliZ* (Bok *et al*., 2006; Ries *et al*., 2020). Also, SM production of BGCs that do not contain a cluster-specific transcription factor but are instead controlled by global regulators, is affected by the loss of these global regulators (Brakhage, 2013). For example, to ensure the production of fumiquinazoline C (Fqm C), a cytotoxic peptidyl alkaloid present in conidia, the coordination of the oxidoreductase FmqD with the conidiation-specific transcription factor BrlA is required (Twumasi-Boateng *et al*., 2009; Lim *et al*., 2014). Loss of *brlA* led to undetectable transcripts of the *fmq*-BGC (Lim *et al*., 2014). Likewise, the MpkA-regulated transcription factor RlmA binds to the promoters of FmqA and FmqC and its loss led to poor production of FqmC (Rocha *et al*., 2021). The significant reduction of *has, hel* and *xan* BGC transcription and production of their encoded compounds in the Δ*smpR* mutant strain indicates that SmpR is not the sole regulator of these BGCs but instead functions as part of a broader multi-pathway regulatory network orchestrating the expression of multiple BGCs in *A. fumigatus*. In agreement with this hypothesis, the involvement of the global regulator LaeA was at least partially demonstrated for the *has* and *hel* BGCs (Perrin *et al*., 2007). Furthermore, it was shown that HAS production is regulated by SreA and HapX in an iron-dependent manner, while AceA and MacA regulate the *xan* BGC under copper repletion (Wiemann *et al*., 2014; Lim *et al*., 2018). Recently, the transcription factors NsdD and Skn7 were established as negative regulators of the *has* and *hel* BGCs, respectively, while RofA may act as a positive regulator of the *hel* BGC (Seo *et al*., 2025). While the cluster-specific transcription factor FccR is ultimately responsible for the regulation of the *fcc* BGC, global regulators for this BGC were unknown (König, 2015). Here, we demonstrated that the production of fumicyclines largely depends on SmpR. Furthermore, we observed that co-cultivation of *S. rapamycinicus* with an overexpressing of *smpR* strain under inducing conditions led to different outcomes in target BGC gene expression. Especially for the *has* BGC, the additional co-culture increased gene expression for up to 40-fold, while this effect was not as pronounced for the *hel, xan* and *fcc* BGCs, suggesting that bacteria*-*associated factors, can further fine-tune the regulation of these BGCs.

Interestingly, the SmpR-regulated BGCs are known for their antimicrobial activity. Helvolic acid is a potent compound against Gram-positive bacteria, especially *Staphylococcus aureus* and methicillin-resistant *S. aureus* (MRSA), although less potent against Gram-negatives (Sang *et al*., 2024; Kamei and Watanabe, 2005). Antibacterial activity against different bacterial species was also reported for the copper-chelator Xanthocillin (Raffa *et al*., 2021). While the antimicrobial activity for HAS, which chelates iron and is discussed as virulence factor of *A. fumigatus,* is not directly evident, it is likely that it can contribute to an antibacterial response of *A. fumigatus* against bacteria by competing for free iron (Yin *et al*., 2013). Fumicyclines were reported to be moderately active against *S. rapamycinicus* on agar plate assays (König *et al*., 2013). Here we could confirm and extend these findings to different soil-dwelling bacteria. Both *Kribbella spp*., and *Arthrobacter* spp., two soil bacteria belonging to the phylum Actinobacteria and known to produce antifungal SMs, activated both SmpR as well as fumicycline production which was not observed with *B. subtilis* and *E. coli* (Evtushenko and Krausova, 2015; Roy and Kumar, 2020; Ramlawi *et al*., 2021; Virués-Segovia *et al*., 2022). It is surprising that the soil isolates did not increase the transcript levels of genes of the *has*, *hel* or *xan* BGCs, suggesting some specificity of the induction by these bacteria which might be linked to the antibacterial activity of especially fumicycline B that inhibited the growth of both soil isolates in concentrations measured in the co-cultivation of these bacteria with *A. fumigatus*.

That the ability to activate fumicycline production is not a universal trait is supported by the finding obtained with several *Streptomyces* spp.. It is linked to the ability of the latter bacteria to produce arginoketides, like AzF, as also shown for the induction of the silent *ors* gene cluster in *A. nidulans* (Krespach *et al*., 2023). While the inducing bacteria *S. iranensis* and *S. rapamycinicus* are closely related and belong to one clade, *S. hawaiiensis* belongs to a different clade within the *Streptomycetaceae* (Labeda *et al*., 2012). It is thus conceivable that *S. hawaiiensis* produces other arginoketides than AzF that induce the fumicycline production, as shown for the production of orsellinic acid in *A. nidulans* (Krespach *et al*., 2023). It is interesting to note that irrespective of the ability of *Streptomyces* spp. to induce the fumicycline production, the compounds are able to reversibly inhibit *Streptomyces* spp. growth, with the strength of this effect depending on the fumicycline isomer.

The expression of SmpR itself needs a specific stimulus, which might derive from the fungal cell membrane because also amphotericin B showed moderate inducing activity. Amphotericin B is known to bind to ergosterol in the fungal cell membrane and induces pores (Mesa-Arango *et al*., 2012). While the mode of action of AzF on fungal cells has not been elucidated yet, it may also target bacterial cell membranes, because Azalomycin F_5a_, a part of the AzF complex, can increase membrane permeability and disrupt the phospholipid bilayer of MRSA (Yuan *et al*., 2019). Its antibacterial activity was shown to be overcome by adding polar lipids to the culture (Krespach *et al*., 2020). It is thus conceivable that *smpR* transcription is activated by compounds that disturb the cell membrane and not generally by SMs since other here tested SMs failed to induce *smpR* transcription.

Taken together, our data suggest that SmpR acts as a hierarchically upstream regulator of FccR and thus the fumicycline BGC. SmpR represents a global regulatory element that also controls the expression of at least the *has, hel* and *xan* BGC. The transcription factor appears to coordinate the antimicrobial response of *A. fumigatus* against *S. rapamycinicus* by regulating several genes discussed as mediators of antibacterial defence. A defence response against bacterial and fungal competitors, also described as the fungal immune system, has been recently described in the basidiomycete *Schizophyllum commune.* It showed some similarities with the *A. fumigatus* response described here, e.g. the activation of several ABC transporter genes (Beijen *et al*., 2024).

## Acknowledgements

We thank Silke Steinbach and Christina Täumer for excellent technical assistance, Moemi Kawashima for providing plasmid pMK024 and Maria Stroe for initial MS measurements. This work was funded by the Federal Ministry of Education and Research (BMBF) within the ANR/BMBF 2019 Antimicrobial resistance call, project titled “Antifungal Resistance: From Surveillance to Treatment” (AReST, grant number: 01KI2120A) to AAB and BH, the Deutsche Forschungsgemeinschaft (DFG) Collaborative Research Center (CRC) 1127 ChemBioSys (Project-ID 239748522) to AAB and the DFG CRC / Transregio 124 FungiNet (INF; 210879364) to GP and SS. AEB was funded by the Cluster of Excellence “Balance of the Microverse – Project ID 390713860)”.

