## supplements for "The fungal transcription factor SmpR coordinates secondary metabolism and antibacterial defence in *Aspergillus fumigatus* during interspecies interaction"

### Generation of mutant strains

**Deletion of *smpR* (Afu1g15910)**

For the generation of the *smpR* deletion mutant (*ΔsmpR)* a 2000 bp DNA sequence homologous to the 5´and 3´ flanking regions of *smpR* was amplified from Δ*KU80* DNA using primer pairs P1/P2 and P5/P6 respectively. The pyrithiamine (*ptrA*) resistance cassette was amplified from plasmid pSK275 (Krappmann *et al.*, 2006) using primer pair P3/P4. The *ptrA* fragment was fused with the gene flanking regions using the Phusion Flash High-Fidelity PCR Master Mix (Thermo Fisher Scientific, Dreieich, Germany) and primer pair P1/P6. The construct was used for transformation of *A. fumigatus* Δ*KU80* protoplasts and transformants were selected on AMM agar plates containing 100 µg/ml pyrithiamine (Sigma-Aldrich, Taufkirchen, Germany).

**Complementation of the Δ*smpR* strain**

For complementation of the *smpR* knockout mutant strain, the gene and its 5´ flanking region were amplified from Δ*KU80* DNA using primer pair P7/P8, the 3´ flanking region was amplified using primer pair P11/P12. The hygromycin (*hph*) resistance cassette and the vector backbone were amplified from plasmid pUC18hph (Liebmann *et al.*, 2004) using primer pairs P9/P10 and P12/P13. All amplifications were carried out using the Phusion Flash High-Fidelity PCR Master Mix (Thermo Fisher Scientific, Dreieich, Germany). The resulting DNA fragments were assembled with Gibson cloning (Gibson, 2009) utilizing the NEBuilder HiFi DNA assembly Master Mix (New England Biolabs, Frankfurt am Main, Germany) following the manufacturer´s instruction. After transformation of *E. coli*, the resulting plasmid pST_002_*smpR*$^{C}$-hph_pUC18 was linearized with *Dra*I (New England Biolabs, Frankfurt am Main, Germany) and used to transform Δ*smpR* protoplasts. Transformants were selected on AMM agar plates containing 150 µg/ml hph (Invivogen, Toulouse, France).

For complementation of the *ΔsmpR* mutant strain with the *smpR*-nanoluciferase (*nluc*)*,* the *smpR* gene and its 3´ flanking region were amplified from Δ*KU80* DNA using primer pair P7/P15, amplification of the 3´gene flanking region was carried out using primer pair P22/P12 using Δ*KU80* DNA. The nanoluciferase gene with an attached linker as well as a terminator (ticl) were amplified from plasmid pMK024 using primer pairs P16/P17 and P18/P19, respectively. The *hph* gene cassette and the vector backbone were amplified from pST_002_*smpR*$^{C}$-hph_pUC18 using primer pairs P20/P21 and P23/P24 respectively. The resulting DNA fragments were assembled with Gibson cloning (Gibson, 2009) utilizing the NEBuilder HiFi DNA assembly Master Mix following the manufacturer´s instruction. After transformation of *E. coli,* the resulting plasmid pST_018_*smpR*-nluc-hph_pUC18 was linearized with *Dra*I. Transformation and selection were carried out as described above.

**Generation of promoter replacement strains**

To generate the $Xyl_{P}$-*fccR, ΔsmpR* construct, the 5´gene flanking region of *fccR* was amplified from Δ*KU80* DNA using primer pair P36/P37. The *hph* resistance cassette was amplified from plasmid pUC18hph (Liebmann *et al.*, 2004) utilizing primer pair P38/P39. The xylose promoter was amplified from pSK529 (Hartmann *et al.*, 2010; Jimenez-Ortigosa *et al.*, 2012) with primer pair P29/P40. Primer pair P41/P42 was used to amplify *fccR* from *ΔKU80* DNA. The vector backbone was amplified from pST_002_$smpR^{C}$-hph_pUC18 utilizing primer pair P43/P44. The resulting DNA fragments were assembled with Gibson cloning (Gibson, 2009) utilizing the NEBuilder HiFi DNA assembly Master Mix (New England Biolabs, Frankfurt am Main, Germany) following the manufacturer´s instruction yielding plasmid pST_005_ hph-$Xyl_{P}$-*fccR*_pUC18. The final construct for $Xyl_{P}$-*fccR, ΔsmpR* was amplified from pST_005_ hph-$Xyl_{P}$-*fccR*_pUC18 utilizing primer pair P36/P42 and was used to transform *ΔsmpR* protoplasts. Transformants were selected on AMM agar plates containing 150 µg/ml hph (Invivogen, Toulouse, France).

For the generation of promoter replacement strains $Xyl_{P}$-*smpR* and $Xyl_{P}$-*smpR, ΔfccR* primer pairs P1/P26 and P5/P6 were used to amplify the 5´ and 3´ gene flanking region from Δ*KU80* DNA respectively. For $Xyl_{P}$-*smpR*, the *ptrA* cassette was amplified from pSK275 (Krappmann *et al.*, 2006) utilizing primer pair P27/P28. For $Xyl_{P}$-*smpR, ΔfccR* the *hph* resistance cassette was amplified from pUC18hph (Liebmann *et al.*, 2004) using primer pair P34/P35. The xylose promoter employed for the generation of both strains was amplified from pSK529 (Hartmann *et al.*, 2010; Jimenez-Ortigosa *et al.*, 2012) with primer pair P29/P30. *smpR* was amplified from Δ*KU80* DNA with primer pair P31/P8. Primer pair P32/P33 was used to amplify the pUC18 vector backbone from pST_002_$smpR^{C}$-hph_pUC18. The resulting DNA fragments for each strain were assembled with Gibson cloning (Gibson, 2009) utilizing the NEBuilder HiFi DNA assembly Master Mix (New England Biolabs, Frankfurt am Main, Germany) following the manufacturer´s instruction. After transformation of *E. coli,* the resulting plasmid pST_006_ptrA-$Xyl_{P}$-*smpR*_pUC18 was linearized with *Dra*I and used to transform *ΔKU80* protoplasts. Plasmid pST_014_hph-$Xyl_{P}$-*smpR*_pUC18 was linearized with *Dra*I and used to transform *ΔfccR* protoplasts. Selection of transformants was carried out using AMM agar plates with 100 µg/ml pyrithiamine (Sigma-Aldrich, Taufkirchen, Germany).

### Verification of strains

Mutants strains were verified by Southern blot analyses. 10 µg of genomic DNA were digested with *Eco*RI (Δ*smpR, smpR-nluc*), *Apa*I ($smpR^{C}$), *Bam*HI ($Xyl_{P}$-*fccR*, Δ*smpR*), *Zra*I ($Xyl_{P}$-*smpR*) or *Xho*I ($Xyl_{P}$-*smpR,* Δ*fccR).*

DNA fragments were separated on an 0.8% (w/v) agarose gel and blotted onto Hybond N+ nylon membranes (GE Healthcare Bio-Sciences, Düsseldorf, Germany). Labelling of DNA probes, hybridization and detection of DNA–DNA hybrids were performed as described before (Grosse *et al.*, 2008). The probes for Southern blot analysis were amplified using primers P5/P6 (Δ*smpR*), P11/P12 (*smpR*$^{C}$), P1/P2 (*smpR-nluc*), P36/P37 (*Xyl*$_{P}$-*fccR*, Δ*smpR*), P5/P6 (*Xyl*$_{P}$-*smpR*), and P1/P26 (*Xyl*$_{P}$-*smpR,* Δ*fccR*).

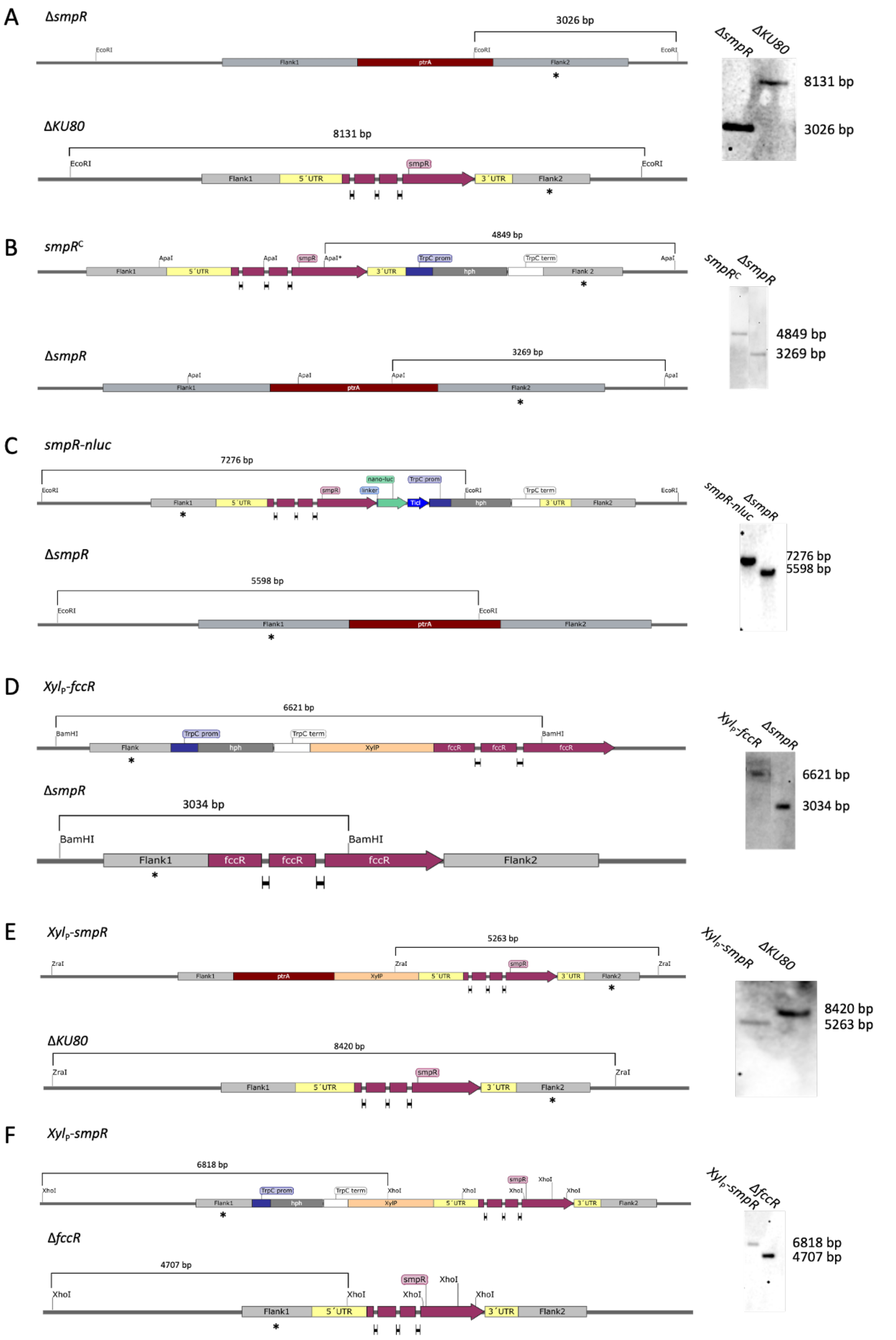


**Figure S1** **Generation of strains used in this study and their verification.** A) Δ*smpR* B) *smpR*$^{C}$ C) *smpR-nluc* D) *Xyl*$_{P}$*-fccR*, Δ*smpR* E) *Xyl*$_{P}$*-smpR* F) *Xyl*$_{P}$*-smpR*, Δ*fccR*. Arrows indicate genes, grey boxes represent 5´and 3´gene flanking regions with 5´ and 3´untranslated regions (UTR) marked in light yellow. Red boxes represent *ptrA* resistance cassettes, dark blue, dark grey and white boxes represent *hph* promoter, *hph* cassette and *hph* terminator, respectively. Light orange boxes represent $Xyl_P$ while the green box marks *nluc*. Flanking regions used as probes for Southern blots are marked with asterisks. *(Right)* Southern blot hybridizations show the expected bands after restriction digest of DNA of mutants and their respective wild types.

### Analysis of gene transcript levels

To verify altered transcript levels of the mutant strains, qRT-PCR analysis was carried out. RNA isolation, DNase digestion, reverse transcription into cDNA and qRT-PCR setup were done as described in Material and Methods. *smpR* transcript was amplified using qPCR primers *smpR*_qP_for/*smpR*_qP_rev; *fccR* transcript was amplified using qPCR primers *fccR*_qP_for/*fccR*_qP_rev. Target gene transcript levels were normalized based on the expression of *act1*, which was amplified with primer pair actin_qP_for/actin_qP_rev, using the $2^{-\Delta\Delta Ct}$ method (Livak and Schmittgen, 2001). The transcript levels of the respective target gene in co-culture with *S. rapamycinicus* were used as control. Statistical analysis was carried out using the GraphPad 10 software (GraphPad Software, Inc, San Diego, CA, USA). A one-way ANOVA with Tukey´s multiple comparisons test was used to compare the means of each experimental group with the means of every other experimental group. Differences between the groups were considered significant at $p \leq 0.05$. Statistical test results were included in the figure with different letters indicating statistical significance, while same letters show non-significance.

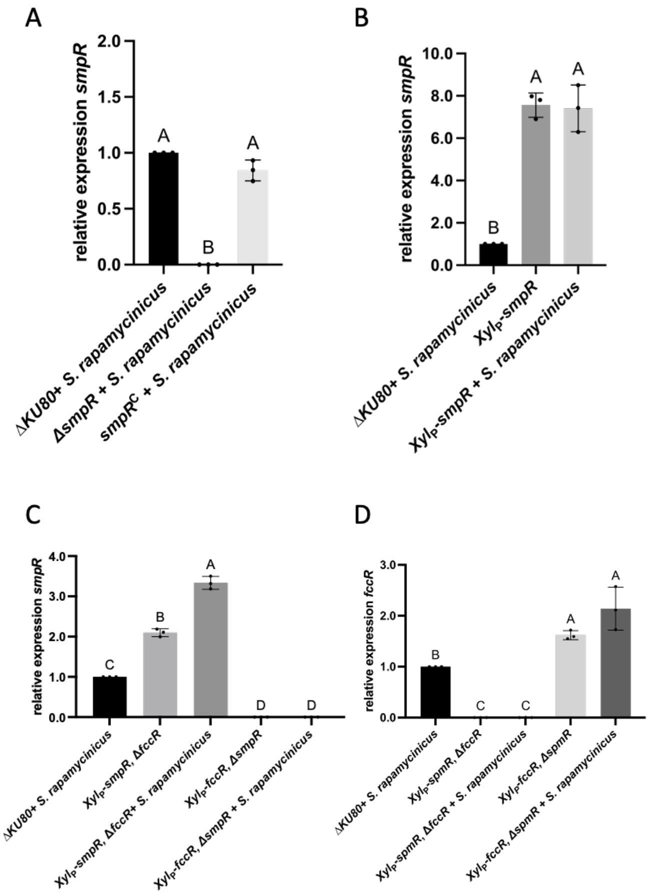


**Figure S2** **Relative expression of** A) *smpR* in Δ*smpR* + *S. rapamycinicus* and *smpR*$^{C}$ + *S. rapamycinicus* B) *smpR* in *Xyl*$_{P}$-*smpR* and *Xyl*$_{P}$-*smpR* + *S. rapamycinicus* C) *smpR* in *Xyl*$_{P}$-*smpR*, Δ*fccR, Xyl*$_{P}$-*smpR*, Δ*fccR* + *S. rapamycinicus*, *Xyl*$_{P}$-*fccR*, Δ*smpR, Xyl*$_{P}$-*fccR*, Δ*smpR* + *S. rapamycinicus* and D) *fccR* in *Xyl*$_{P}$-*smpR*, Δ*fccR, Xyl*$_{P}$-*smpR*, Δ*fccR* + *S. rapamycinicus*, *Xyl*$_{P}$-*fccR*, Δ*smpR, Xyl*$_{P}$-*fccR*, Δ*smpR* + *S. rapamycinicus*

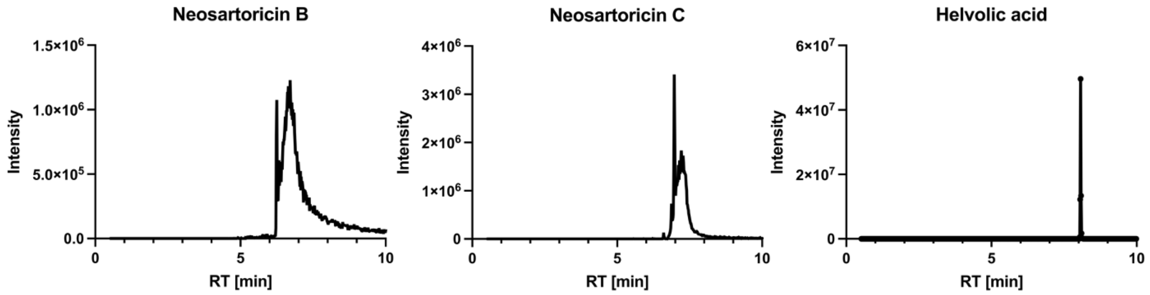


**Figure S3** **LC-MS analysis.** EICs of neosartoricin B, neosartoricin C and helvolic acid standards.

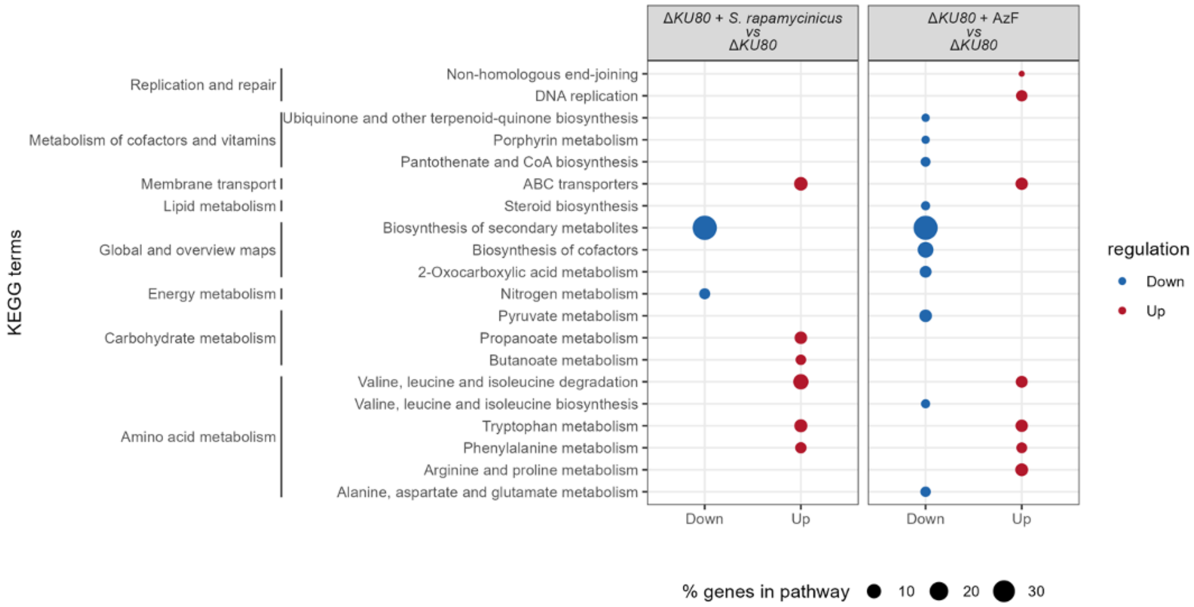


**Figure S4 KEGG terms of *A. fumigatus* Δ*KU80* wild typeco-cultured with *S. rapamycinicus* or treated with AzF.** KEGG terms are depicted according to their abundance. Lower transcribed terms are labelled in blue, higher transcribed terms in red.

### Determination of Minimal Inhibitory Concentration (MIC) and Minimal Effective Concentration (MEC)

MICs and MEC were determined by the EUCAST broth dilution methodology of antifungal agents for conidia-forming moulds (EUCAST, 2020). Briefly, stock solutions of azalomycin F (Maira Rosin, Leibniz-HKI), amphotericin B (Sigma-Aldrich, Taufkirchen, Germany), voriconazole (Sigma-Aldrich, Taufkirchen, Germany) and caspofungin (Biomol, Hamburg, Germany) were dissolved in dimethyl sulfoxide (DMSO). All agents were tested over a range of 0.0625 µg/ml - 32 µg/ml. MICs were defined as the lowest concentration with microscopically observable complete inhibition of fungal growth after 48 h of incubation in AMM (with or without xylose where required). The MEC for caspofungin was defined as the lowest concentration with microscopically observable effect on fungal morphology after 48 h of incubation (Table S9).

### Characterization of *A. fumigatus* Δ*smpR* deletion strain

#### Spot assays

For the assessment of fungal growth on minimal (AMM) agar plates or rich media (Malt agar), conidia of strains Δ*smpR, smpR*$^C$, *Xyl*$_P$-*smpR* and Δ*KU80* wild type, were diluted in ultrafiltrated water to a concentration of $10^5$/ml. Five microliter were spotted on either AMM agar or Malt agar. Colony diameters were documented after incubation for 48 h at 37 °C.

For the assessment of fungal growth under different stress conditions, freshly harvested conidia were serially diluted in double distilled water to concentrations of $10^5$, $10^4$, $10^3$, $10^2$/ml. Five microliter of conidial dilutions were spotted on AMM agar plates in the presence of stress-inducing agents, as indicated. Colony diameters were documented as indicated. To induce the overexpression of *smpR*, conidia were spotted on xylose-containing AMM agar plates.

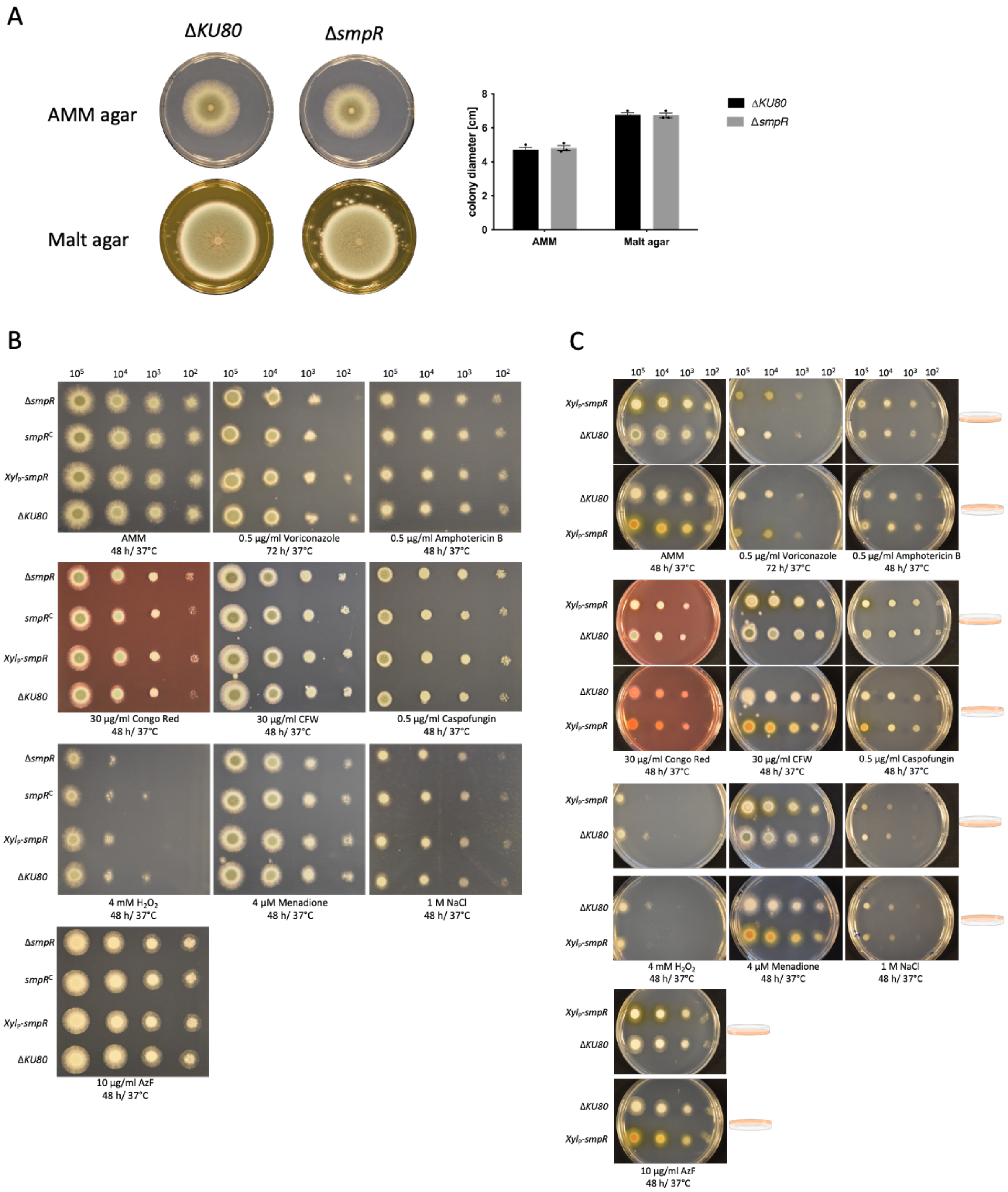


**Figure S5** **Growth of strains Δ*smpR*, *smpR*$^{C}$, *Xyl*$_{P}$*-smpR* and Δ*KU80*** A) Growth of Δ*KU80* and Δ*smpR* on AMM agar and Malt agar; *(Right)* analysis of colony diameter of Δ*KU80* and Δ*smpR*. B) Growth of Δ*smpR*, *smpR*$^{C}$,

*Xyl*P-*smpR* and Δ*KU80* on Glucose-AMM agar containing stress-inducing agents. C) Growth of *Xyl*P-*smpR* and Δ*KU80* on Xylose-AMM agar containing stress-inducing agents.

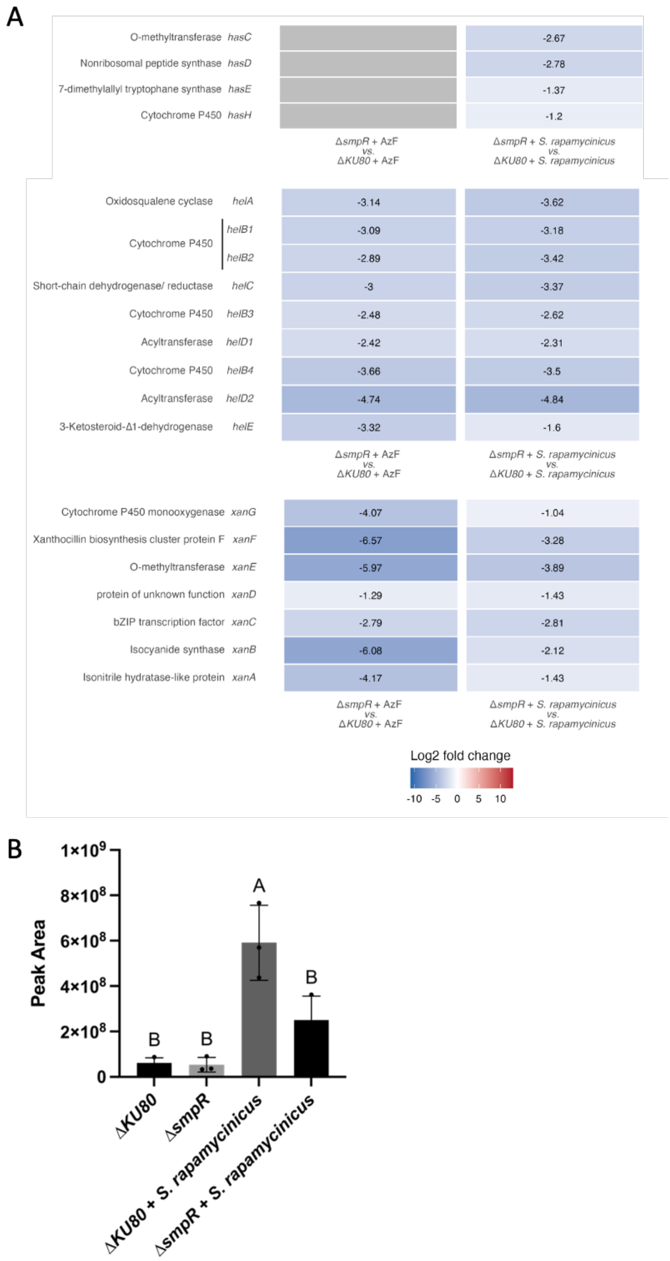


**Figure S6** **Analysis of transcript levels of the *has, hel* and *xan* BGC.** A) Transcript levels of the *top has, middle hel* and *bottom xan* BGCs in Δ*KU80* wild type and Δ*smpR* mutant strain. Genes are displayed with name and $\log_2$ fold changes during AzF treatment or *S. rapamycinicus* co-culture. The scale used defines the expression of genes from lower expressed (blue) to higher expressed (red). Genes without significant fold change are displayed in white. Grey bars indicate genes that were not found in the analysis. B) Helvolic acid production in Δ*KU80* and Δ*smpR* in axenic culture or co-culture with *S. rapamycinicus*.

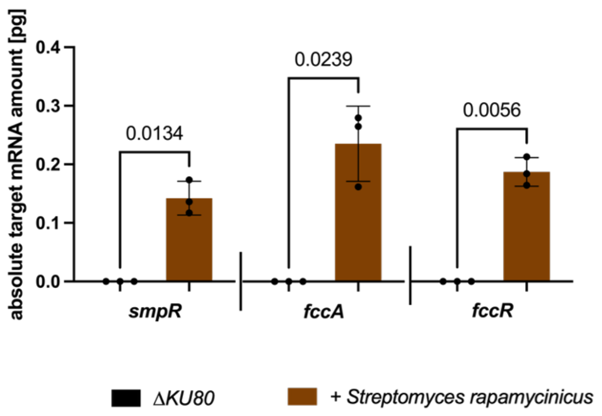


**Figure S7** **Analysis of transcript levels of genes *smpR, fccA* and *fccR* of axenic or co-cultured *A. fumigatus* Δ*KU80* wild type**. Transcript levels of *smpR, fccA* and *fccR* were determined *via* qRT-PCR as absolute mRNA quantification.

### Supplementary Tables

**Table S1.** ***Aspergillus fumigatus* strains used in this study**

| Strains | Genotype | Source |
|---|---|---|
| Δ*KU80* | *KU80Δ* | da Silva Ferreira *et al.*, 2006 |
| Δ*smpR* | *KU80Δ; Afu1g15910::ptrA* | This study |
| *smpR*$^C$ | *KU80Δ; Afu1g15910::ptrA*; *smpR-hph* | This study |
| *smpR*-nluc | *KU80Δ; Afu1g15910::ptrA; smpR-nluc-hph* | This study |
| *XylP-smpR* | *KU80Δ; ptrA-Xyl*$_P$*-smpR* | This study |
| *Xyl*$_P$*-smpR, ΔfccR* | *KU80Δ; hph-Xyl*$_P$*-smpR; fccR::ptrA* | This study |
| *Xyl*$_P$*-fccR, ΔsmpR* | *KU80Δ; hph-Xyl*$_P$*-fccR; smpR0::ptrA* | This study |

**Table S2.** ***Streptomyces* strains used in this study**

| Strain | Genotype | Source |
|---|---|---|
| *Streptomyces rapamycinicus* ATCC 29253 | Wild-type strain | Kumar and Goodfellow, 2008 |
| *Streptomyces iranensis* JCM 17327 | Wild-type strain | Hamedi et al., 2010 |
| *Streptomyces coelicolor* A3(2) | Wild-type strain | Erikson, 1955 |
| *Streptomyces lividans* PM02 | Wild-type strain | van Dissel *et al.*, 2015 |
| *Streptomyces fradiae* DSM10745 | Wild-type strain | Waksman and Curtis, 1916 |
| *Streptomyces hawaiiensis* ATCC 12236 | Wild-type strain | Cron *et al.*, 1956 |
| *Streptomyces viridifaciens* ATCC 31495 | Wild-type strain | Gourevitch and Lein, 1955 |
| *Streptomyces nodosus* ATCC 14899 | Wild-type strain | Trejo and Bennett, 1963 |
| *Streptomyces griseus* JCM 4626 | Wild-type strain | Waksman *et al.*, 1948 |

**Table S3.** **Bacterial strains used in this study.**

| Strain | Genotype | Source |
|---|---|---|
| *Bacillus subtilis* 168 | Wild-type strain | Nakamura *et al.*, 1999 |
| *Escherichia coli* α-Select chemically competent cells | | Bioline, GmbH, Luckenwalde, Germany |
| *Arthrobacter* spp. | Wild-type strain | This study |
| *Kribbella* spp. | Wild-type strain | This study |

**Table S4.** **Primers used in this study**

| Name | Sequence (5´-3´) |
|---|---|
| OMK_505 | AGAGTTTGATCMTGGCTCAG |
| OMK_506 | TACGGYTACCTTGTTACGACTT |
| P1_*ΔsmpR*_Flank1_for | CCCGACTGAACATGGTAGGT |
| P2_*ΔsmpR*_Flank1_rev | TATGTCTGGTGATAATTGGG |
| P3_*ΔsmpR*_*ptrA*_for | TAACCGCTACCCCAATTATCACCAGACATAatggcctagatggcctcttg |
| P4_*ΔsmpR*_*ptrA*_rev | GGATTTGAGAAGTGAAAAGTGAGAGTGCGGtcaggccaattgattacggg |
| P5_*ΔsmpR*_Flank2_for | CCGCACTCTCACTTTTCACT |
| P6_*ΔsmpR*_Flank2_rev | CTCGATCAGGGAGGTGAAGA |
| P7_*smpR*$^C$_Flank1 *smpR*_for | CCCGACTGAACATGGTAGGT |
| P8_*smpR*$^C$_Flank1 *smpR*_rev | GCGGGCCGGGATTGGGGAAT |
| P9_*smpR*$^C$_hph_for | ATTCCCCAATCCCGGCCCGCgacgttaactgatattgaag |
| P10_*smpR*$^C$_hph_for | AAAATATCCATCCACCACGTaacccaggggctggtgacgg |
| P11_*smpR*$^C$_Flank2_for | ACGTGGTGGATGGATATTTT |
| P12_*smpR*$^C$_Flank2_rev | TTGTTGCAGCACATGATCAC |
| P13_*smpR*$^C$_Vector_rev | GTGATCATGTGCTGCAACAATTTAAAgggtaccgagctcgaattcg |
| P14_*smpR*$^C$_Vector_rev | ACCTACCATGTTCAGTCGGGTTTAAAggggatcctctagagtcgac |
| P15_Nluc_Flank1 *smpR*_rev | GGTGCAGATAGTTCCATTCC |
| P16_Nluc_for | GGAATGGAACTATCTGCACCGGTGGAGGCTCTatggtctt |
| P17_Nluc_rev | GCTCTTGACTATCCACCATCTCAcgccagaatgcgttcgcaca |
| P18_Nluc_ticl_for | GATGGTGGATAGTCAAGAGC |
| P19_Nluc_ticl_rev | cttcaatatcagttaacgtcCTGCTAAGACCGCAGTAGTC |
| P20_Nluc_hph_for | gacgttaactgatattgaag |
| P21_Nluc_hph_rev | agtgaaaagtgagagtgcggaacccaggggctggtgacgg |
| P22_Nluc_Flank2_for | ccgcactctcacttttcact |
| P23_Nluc_vector_for | GTGATCATGTGCTGCAACAATTTAAAgggtaccgagctcgaattcg |
| P24_Nluc_vector_rev | ACCTACCATGTTCAGTCGGGTTTAAAggggatcctctaga |
| P26_*Xyl*$_P$-*smpR*_Flank1_rev | GATTGATCATTGATGTATGG |
| P27_*Xyl*$_P$-*smpR*_ptrA_for | CCATACATCAATGATCAATCatggcctagatggcctcttg |
| P28_*Xyl*$_P$-*smpR*_ptrA_rev | ccgtCGGCGGCCGCCTGCAGtcaggccaattgattacggg |
| P29_*Xyl*$_P$-*smpR*_Xyl$_P$_for | CTGCAGGCGGCCGCCGacgg |
| P30_*Xyl*$_P$-*smpR*_Xyl$_P$_rev | GTCAGATGCTGATCGTTTGGgttggttcttcgagtcgatg |
| P31_*Xyl*$_P$-*smpR*_*fccM*_for | CCAAACGATCAGCATCTGAC |
| P32_*Xyl*$_P$-*smpR*_Vector_for | ATTCCCCAATCCCGGCCCGCTTTAAAgggtaccgagctcgaattcg |
| P33_*Xyl*$_P$-*smpR*_Vector_rev | ACCTACCATGTTCAGTCGGGTTTAAAggggatcctctagagtcgac |
| P34_ *Xyl*$_P$-*smpR*_hph_for | CCATACATCAATGATCAATCgacgttaactgatattgaag |
| P35_ *Xyl*$_P$-*smpR*_hph_rev | ccgtCGGCGGCCGCCTGCAGaacccaggggctggtgacgg |
| P36_*Xyl*$_P$-*fccR*_Flank1_for | GAGATGATTGGCGAGGCAGA |
| P37_*Xyl*$_P$-*fccR*_Flank1_rev | CTTGATAATAGATGGTTGTT |
| P38_*Xyl*$_P$-*fccR*_hph_for | AACAACCATCTATTATCAAGgacgttaactgatattgaag |
| P39_*Xyl*$_P$-*fccR*_hph_rev | ccgtCGGCGGCCGCCTGCAGaacccaggggctggtgacgg |
| P40_*Xyl*$_P$-*fccR*_Xyl$_P$_rev | TCCGCCGATTACTTTTCTCCATgttggttcttcgagtcgatg |
| P41_*Xyl*$_P$-*fccR*_*fccR*_for | ATGGAGAAAAGTAATCGGCGGA |
| P42_*Xyl*$_P$-*fccR*_*fccR*_rev | CTAAGGAAGAAGCTGGTTCGAC |
| P43_*Xyl*$_P$-*fccR*_Vector_for | CGAACCAGCTTCTTCCTTAGgggtaccgagctcgaattcg |
| P44_*Xyl*$_P$-*fccR*_Vector_rev | TCTGCCTCGCCAATCATCTCTTATAAggggatcctctaga |

**Table S5.** **qPCR primers and their efficiency used in this study**

| Name | Sequence (5´-3´) | Efficiency [%] |
|---|---|---|
| *fccA*_qP_for | ATCAGAACCCGGACGCGAAC | 95.3 |
| *fccA*_qP_rev | GGGATTGTCCAGCCAGGAGC | |
| *fccB*_qP_for | TGCTTCTTACTTGAGGAGGAG | 91.64 |
| *fccB*_qP_rev | TGTGCGGGGTATCCAATCAC | |
| *fccC*_qP_for | GCCGTGGAGTCTGTTTCTCAATG | 94.7 |
| *fccC*_qP_rev | CGCCAAATATGGGCATCGCAT | |
| *fccD*_qP_for | TGTGGTATGAGCTGAACGGA | 104.4 |
| *fccD*_qP_rev | TTGACATCAGGAAGAGCAAGA | |

| | | |
|---|---|---|
| *fccE*_qP_for<br>*fccE*_qP_rev | CGTATGGGATGCTGGCCAAAG<br>ACAGCCTTCCCATTCGAAGCC | 105.4 |
| *fccR*_qP_for<br>*fccR*_qP_rev | ATTGCCAGAAGGTCAGAAGAC<br>TGGTGAGCCACAACCGC | 107.71 |
| *smpR*_qp_for<br>*smpR*_qP_rev | TGTCAGTCGGACCTGTGGA<br>CCATTGAAGACGGCTTTTG | 98.8 |
| actin_qP_for<br>actin_qP_rev | GGGATGACATGGAGAAGATCTG<br>GCGTTGAAGGTCTCGAAGAC | 104.2 |
| *hasC*_qP_for<br>*hasC*_qP_rev | AGTATCTGGCCGAAACTGGCA<br>CCGACTTGACTTCCTCCCATGGG | 96.5 |
| *hasD*_qP_for<br>*hasD*_qP_rev | CCTGGGTGCAGAATCGCTGT<br>GCAACCTTGGAACCCACAGACTAT | 98.7 |
| *hasE*_qP_for<br>*hasE*_qP_rev | CCGCACGTATATCTTGATCCGA<br>AGCGCTGAAGAGTGAGGTGT | 91.5 |
| *hasH*_qP_for<br>*hash*_qP_rev | ATGCCGGCTGGCTCTTGACT<br>GGCTGTCGCGCAAATACTGC | 100.7 |
| *helA*_qP_for<br>*helA*_qP_rev | CTTCTCGCATCTCCGATGAGC<br>CGGTGATGTTGAGCAGTTCCAG | 95.6 |
| *helB1*_qP_for<br>*helB1*_qP_rev | TTGCCGTATGTCGAAGCCTG<br>GATTTCTGTCCCTGGCGGAATG | 101.5 |
| *helB2*_qP_for<br>*helB2*_qP_rev | CAACGGGCTTCTGGCATCCT<br>GCGTGCGGATCAATTTCCGGTT | 94.6 |
| *helC*_qP_for<br>*helC*_qP_rev | GCCACCCAACAGGCTATCAAGTAC<br>CCCCAGCCGCAAAGGTAAACA | 94.2 |
| *helB3*_qP_for<br>*helB3*_qP_rev | GAATCTGGATCTTCCGCGTCGA<br>CCTAGTGGCCTTTAGCCAGGAG | 96.7 |
| *helD1*_qP_for<br>*helD1*_qP_rev | AGCTCGCTCCGTAGGTAAGTAC<br>CTTTTATCAATCACTGGGCGGAG | 99.6 |
| *helB4*_qP_for<br>*helB4*_qP_rev | GGTGAGAGGGAGAGAACTGGCT<br>TGAGGTGATGACAGAGAGCATTG | 101.7 |
| *helD2*_qP_for<br>*helD2*_qP_rev | GCAGGATGCCACATCGAGCG<br>TTGGGGATGGTTTGCCAGGAACC | 103.6 |
| *xanA*_qP_for<br>*xanA*_qP_rev | GCTCGATCACGAACAGTGAACG<br>GTGGCCCACCAGGGACTACCAT | 99.8 |
| *xanB*_qP_for<br>*xanB*_qP_rev | GGGAGCCACAAGAGCATAACCTA<br>CAGCGAGGTCTTCAGGAGCT | 106.8 |
| *xanC*_qP_for<br>*xanC*_qP_rev | GGATATGGCGGGAAGCTGAATTC<br>CTCCACCAACTCATCACCCTCTA | 95.2 |
| *xanD*_qP_for<br>*xanD*_qP_rev | CTCGCTCTCCAGAAACGCGTC<br>TATGCTGTAAGGCGGCCGAGT | 95.1 |
| *xanE*_qP_for<br>*xanE*_qP_rev | GGCGCAAGGAACTGCAACGA<br>AGCGAATCATGCGTGTTTGCAC | 91.7 |
| *xanF*_qP_for<br>*xanF*_qP_rev | CTCGTGCAGCTCAGCAAAGG<br>TGCCCTGTACCAGAGAATCACCG | 96.5 |
| *xanG*_qP_for<br>*xanG*_qP_rev | CGGAACGAGGACCTCTACGCAAA<br>CCCCGTGTACAGCTTCCAGGTGT | 90.2 |

| | | |
|---|---|---|
| *helE*_qP_for | TGCCATCGATTTGACCATGAAACC | 103.6 |
| *helE*_qP_rev | CACCTATTTGGGCATCTTTCTCGA | |
| *aflT*_qP_for | GCCATCATCTGCCTCTTGCT | 102.6 |
| *aflT*_qP_rev | CTGCGAGAATGCAAAGAGGAT | |
| *defX*_qP_for | TCTGCGTGCGCCAGGGTAA | 105.8 |
| *defX*_qP_rev | GGGCTCGTCGGCATCAATCA | |
| AFUA_8G00710_qP_for | TCAGCCAGTGCTACCATCGTC | 103.6 |
| AFUA_8G00710_qP_rev | GGTGGCCTTTTGGAAGTCGAG | |
| AFUA_8G07100_qP_for | CACCACCGGCTATGTCACCAAG | 102.8 |
| AFUA_8G07100_qP_rev | TTCCTTCTTGGCCTCCCCGA | |
| AFUA_8G06980_qP_for | GGATATCGAGTACAACCCCTACGG | 106.3 |
| AFUA_8G06980_qP_rev | GTCTGCAGTGGTGTAGATCATGG | |

**Table S6.** **Plasmids used in this study**

| Plasmid | Source or reference |
|---|---|
| pUC18hph | Liebmann *et al.*, 2004 |
| pMK024 | Moemi Kawashima, personal communication |
| pSK275 | Krappmann *et al.*, 2006 |
| pSK529 | Hartmann *et al.*, 2010; Jimenez-Ortigosa *et al.*, 2012 |
| pST_002_$smpR^{C}$-hph_pUC18 | This study |
| pST_005_ hph-$Xyl_P$-*fccR*_pUC18 | This study |
| pST_006_ptrA-$Xyl_P$-*smpR*_pUC18 | This study |
| pST_014_hph-$Xyl_P$-*smpR*_pUC18 | This study |
| pST_018_*smpR*-nluc-hph_pUC18 | This study |

All plasmids used in this study have been propagated in *E. coli* α-Select. *E. coli* α-Select was grown in LB medium (Carl Roth, Karlsruhe, Germany) supplemented with 60 µg/ml carbenicillin.

**Table S7** **Annotations of genes of Δ*KU80* with differential expression due to treatment with AzF or co-cultivation with *S. rapamycinicus***

| Gene | Annotation |
|---|---|
| Genes with lower transcript levels in co-culture but higher expression during AzF treatment | |
| AFUA_6G11850 | Protein of unknown function |
| AFUA_5G13160 | Has domain(s) with predicted role in transmembrane transport and integral component of membrane localization |
| AFUA_5G13180 | Putative agmatinase, PhoB-regulated |
| AFUA_8G00520 | Fumagillin biosynthesis terpene cyclase |
| AFUA_8G00530 | Alpha/beta hydrolase PsoB |
| AFUA_8G00500 | Putative acetate-CoA ligase |
| AFUA_8G00480 | Nonheme iron-dependent dioxygenase |
| AFUA_8G00560 | Cytochrome P450 monooxygenase PsoD |
| AFUA_1G04870 | Ortholog(s) have putrescine transmembrane transporter activity, spermidine transmembrane transporter activity, urea transmembrane transporter activity and role in putrescine transport, spermidine transport, urea transport |
| AFUA_5G13170 | Putative MATE efflux family protein |
| AFUA_8G00580 | Glutathione S-transferase PsoE |
| AFUA_8G00490 | Stereoselective keto-reductase |
| AFUA_6G11840 | Ortholog(s) have role in calcium ion import and plasma membrane localization |
| AFUA_8G00540 | PKS-NRPS hybrid synthetase PsoA |
| AFUA_8G00440 | Dual-functional monooxygenase/methyltransferase PsoF |
| AFUA_8G00410 | Methionine aminopeptidase type II |
| Genes with higher transcript levels in co-culture but lower expression during AzF treatment | |
| AFUA_5G09900 | Has domain(s) with predicted role in response to stress and integral component of membrane localization |

| | |
|---|---|
| AFUA_4G01140 | Putative multidrug resistance protein |
| AFUA_4G07000 | Atypical dual-specificity phosphatase Siw14-like, putative |
| AFUA_2G10230 | Putative inositol oxygenase |
| AFUA_2G00210 | Ortholog of *A. fumigatus* Af293 : Afu7g06340, *A. niger* CBS 513.88 : An01g11590, An01g01670, *A. oryzae* RIB40 : AO090009000035 and *Aspergillus wentii*: Aspwe1_0027365, Aspwe1_0046529 |
| AFUA_4G09850 | Ortholog of *A. nidulans* FGSC A4 : AN3507, *A. oryzae* RIB40 : AO090103000215, *Neosartorya fischeri* NRRL 181 : NFIA_106200 and *Aspergillus versicolor*: Aspve1_0085394 |
| AFUA_3G14250 | Ortholog(s) have indoleamine 2,3-dioxygenase activity and role in filamentous growth of a population of unicellular organisms in response to chemical stimulus, tryptophan catabolic process to kynurenine |
| AFUA_4G13510 | isocitrate lyase |
| AFUA_4G06570 | Ortholog(s) have hyphal tip localization |
| AFUA_5G09910 | Putative p-nitroreductase family protein |
| AFUA_2G05750 | Ortholog(s) have endoplasmic reticulum, fungal-type vacuole localization |

**Table S8 Annotations of BGCs for fusarinine C, hexadehydroastechrome (HAS), trypacidin, helvolic acid, xanthocillin, pyripyropene, fumicycline and fumagillin/pseurotin A**

| Gene | Annotation |
|---|---|
| | Fusarine C BGC |
| AFUA_3G03390 (*sidJ)* | Siderophore biosynthesis lipase/esterase, putative |
| AFUA_3G03400 (*sidF*) | Siderophore biosynthesis acetylase AceI, putative |
| AFUA_3G03410 (*sidH*) | Enoyl-CoA hydratase/isomerase family protein |
| AFUA_3G03420 (*sidD*) | Nonribosomal peptide synthetase |
| AFUA_3G03430 (*sitT*) | ABC multidrug transporter, putative |
| AFUA_3G03440 (*mirD*) | Putative siderophore transporter |
| | Hexadehydroastechrome BGC |
| AFUA_3G12890 (*hasA*) | C6 transcription factor |
| AFUA_3G12900 (*hasB*) | Putative transporter |
| AFUA_3G12910 (*hasC*) | Putative O-methyltransferase |
| AFUA_3G12920 (*hasD*) | Nonribosomal peptide synthase (NRPS) |
| AFUA_3G12930 (*hasE*) | 7-Dimethylallyl tryptophan synthase |
| AFUA_3G12940 (*hasF*) | C6 transcription factor |
| AFUA_3G12950 (*hasG*) | FAD-binding domain protein |
| AFUA_3G12960 (*hasH*) | Putative cytochrome P450 |
| | Trypacidin BGC |
| AFUA_4G14460 (*tpcM*) | Methyltransferase |
| AFUA_4G14470 (*tpcK*) | Probable decarboxylase |
| AFUA_4G14480 (*tpcL*) | Emodin anthrone oxidase |
| AFUA_4G14490 (*tpcJ*) | Putative dihydrogeodin oxidase |
| AFUA_4G14500 (*tpcI*) | Questin oxygenase, putative |
| AFUA_4G14510 (*tpcH*) | Methyltransferase |
| AFUA_4G14520 (*tpcG*) | Monooqygenase |
| AFUA_4G14530 (*tpcF*) | Putative theta class glutathione s-transferase |
| AFUA_4G14540 (*tpcE*) | Transcription factor |
| AFUA_4G14550 (*tpcD*) | Transcriptional coactivator |
| AFUA_4G14560 (*tpcC*) | Non-reducing polyketide synthase |
| AFUA_4G14570 (*tpcB*) | Trypacidin synthesis protein B |
| AFUA_4G14580 (*tpcA*) | O-Methyltransferase |
| | Helvolic acid BGC |
| AFUA_4G14770 (*helA*) | Oxidosqualene cyclase |
| AFUA_4G14780 (*helB1*) | Cytochrome P450 |
| AFUA_4G14790 (*helB2*) | Cytochrome P450 |
| AFUA_4G14800 (*helC*) | Short-chain dehydrogenase/ reductase |
| AFUA_4G14810 (*helB3*) | Cytochrome P450 |
| AFUA_4G14820 (*helD1*) | Acyltransferase |
| AFUA_4G14830 (*helB4*) | Cytochrome P450 |
| AFUA_4G14840 (*helD2*) | Acyltransferase |
| AFUA_4G14850 (*helE*) | 3-Ketosteroid-$\Delta^1$-dehydrogenase |
| | Xanthocillin BGC |
| AFUA_5G02620 (*xanG*) | Cytochrome P450 monooxygenase |
| AFUA_5G02630 (*xanF*) | Xanthocillin biosynthesis cluster protein F |
| AFUA_5G02640 (*xanE*) | O-methyltransferase |

| | |
|---|---|
| AFUA_5G02650 (*xanD*) | protein of unknown function |
| AFUA_5G02655 (*xanC*) | transcription factor |
| AFUA_5G02660 (*xanB*) | Isocyanide synthase |
| AFUA_5G02670 (*xanA*) | Isonitrile hydratase-like protein |
| | Pyripyropene BGC |
| AFUA_6G13920 (*pyr1*) | Nicotinic acid-CoA ligase |
| AFUA_6G13930 (*pyr2*) | Non-reducing polyketide synthase |
| AFUA_6G13945 (*pyr9*) | Cytochrome P450 monooxygenase |
| AFUA_6G13950 (*pyr4*) | Terpene cyclase |
| AFUA_6G13970 (*pyr5*) | FAD-dependent monooxygenase |
| AFUA_6G13980 (*pyr6*) | Polyprenyl transferase |
| AFUA_6G13990 (*pyr7*) | O-Acetyltransferase |
| AFUA_6G14000 (*pyr8)* | Acetyltransferase |
| | Fumicycline BGC |
| AFUA_7G00120 (*fccB)* | β-Lactamase like thioesterase |
| AFUA_7G00130 (*fccR*) | C6 transcription factor |
| AFUA_7G00150 (*fccC*) | Monooxygenase |
| AFUA_7G00160 (*fccA*) | Polyketide synthase |
| AFUA_7G00170 (*fccD*) | Tryptophan synthase |
| AFUA_7G00180 (*fccE*) | Epimerase/ Dehydratase |
| | Fumagillin/Pseurotin A supercluster |
| AFUA_8G00370 (*fmaB*) | Fumagillin biosynthesis polyketide synthase |
| AFUA_8G00380 (*fmaC*) | Fumagillin biosynthesis acyltransferase |
| AFUA_8G00390 (*fmaD*) | Fumagillin biosynthesis methyltransferase |
| AFUA_8G00400 | Fumagillin biosynthesis methyltransferase |
| AFUA_8G00410 (*fpaII*) | Methionine aminopeptidase type II |
| AFUA_8G00420 (*fapR*) | C6 finger transcription factor |
| AFUA_8G00430 | hypothetical protein |
| AFUA_8G00440 (*psoF*) | Dual-functional monooxygenase/methyltransferase |
| AFUA_8G00450 (*psoG*) | |
| AFUA_8G00460 (*fpaI*) | Methionine aminopeptidase type I, putative |
| AFUA_8g00470 (*fmaE*) | Antibiotic Biosynthesis Monoxygenase superfamily monooxygenase |
| AFUA_8G00480 (*fmaF*) | Nonheme iron-dependent dioxygenase |
| AFUA_8G00490 | Stereoselective keto-reductase |
| AFUA_8G00500 | Putative acetate-CoA ligase |
| AFUA_8G00510 (*fmaG*) | Fumagillin biosynthesis cluster P450 monooxygenase |
| AFUA_8G00520 (*fmaA*) | Fumagillin biosynthesis terpene cyclase |
| AFUA_8G00530 (*psoB*) | Alpha/beta hydrolase |
| AFUA_8G00540 (*psoA*) | PKS-NRPS hybrid synthetase |
| AFUA_8G00550 (*psoC*) | Methyltransferase |
| AFUA_8G00560 (*psoD*) | Cytochrome P450 monooxygenase |
| AFUA_8G00570 | Putative hydrolase |
| AFUA_8G00580 (*psoE*) | Glutathione S-transferase |

**Table S9** **Azalomycin F and amphotericin B MICs of *A. fumigatus* strains Δ*smpR,* Δ*fccR* and Δ*KU80* wild type**

| Strain | Azalomycin F [µg/ml] | Amphotericin B [µg/ml] | Voriconazole [µg/ml] | Caspofungin [µg/ml] |
|---|---|---|---|---|
| Δ*smpR* | 32 | 1 | 1 | 0.5 |
| $smpR^C$ | 32 | 1 | 1 | 0.5 |
| $Xyl_P$-*smpR* | | | | |
| In Glucose AMM | 32 | 1 | 1 | 0.5 |
| In Xylose AMM | 32 | 1 | 0.5 | 1 |
| Δ*KU80* wild type | | | | |
| In Glucose AMM | 32 | 1 | 1 | 0.5 |
| In Xylose AMM | 32 | 1 | 0.5 | 1 |
